# Conserved and cell type-specific transcriptional responses to IFN-γ in the ventral midbrain

**DOI:** 10.1101/2022.12.14.520294

**Authors:** Benjamin D. Hobson, Adrien T. Stanley, Mark B. De Los Santos, Bruce Culbertson, Eugene V. Mosharov, Peter A. Sims, David Sulzer

## Abstract

Dysregulated inflammation within the central nervous system (CNS) contributes to neuropathology in infectious, autoimmune, and neurodegenerative disease. With the exception of microglia, major histocompatibility complex (MHC) proteins are virtually undetectable in the mature, healthy central nervous system (CNS). Neurons have generally been considered incapable of antigen presentation, and although interferon gamma (IFN-γ) can elicit neuronal MHC class I (MHC-I) expression and antigen presentation *in vitro*, it remains unclear whether similar responses occur *in vivo*. Here we directly injected IFN-γ into the ventral midbrain of mature mice and analyzed gene expression profiles of specific CNS cell types. We find that IFN-γ induces cellular proliferation and expression of MHC-II and associated genes only in microglia. However, IFN-γ upregulated MHC-I and associated mRNAs in ventral midbrain microglia, astrocytes, oligodendrocytes, and GABAergic, glutamatergic, and dopaminergic neurons. The core set of IFN-γ-induced genes and their response kinetics were conserved across neurons and glia, with a lower amplitude of expression in neurons. A diverse repertoire of genes was upregulated in glia, particularly microglia, while no neuron-specific responses to IFN-γ were observed. Using mutant mice to selectively delete the IFN-γ-binding domain of IFNGR1 in dopaminergic neurons, we demonstrate that dopaminergic neurons respond directly to IFN-γ. Our results suggest that most neurons are capable of responding directly to IFN-γ and upregulating MHC-I and related genes *in vivo*, but their expression amplitude and repertoire is limited compared to oligodendrocytes, astrocytes, and microglia.

**One-sentence summary:** We find that IFN-γ induces transcription of MHC class I antigen processing and presentation machinery in all major parenchymal cell types in the ventral midbrain; however, neuronal responses are low amplitude and limited to a small set of genes, MHC class II expression and cellular proliferation are restricted to microglia, and dopamine neuronal responses require cell autonomous expression of IFNGR1.

## Introduction

Neuroinflammation is associated with neurodegenerative diseases including multiple sclerosis (MS), Alzheimer’s disease (AD), and Parkinson’s disease (PD) (Amor et al., 2014), while clonally expanded T cells accumulate in the aged brain (Dulken et al., 2019; Gate et al., 2020). MS is a demyelinating disorder classically characterized by autoimmune responses to oligodendrocyte proteins, and recent evidence suggests that T cell reactivity towards the neuronal protein β-synuclein contributes to grey matter pathology in MS (Lodygin et al., 2019). There is considerable neuropathological evidence of inflammation in the substantia nigra pars compacta (SNc) of PD patients, including reactive, HLA-DR^+^ microglia (McGeer et al., 1988) and the presence of CD8+ and CD4+ T cells near the pigmented SNc dopaminergic (DA) neurons and noradrenergic neurons of the locus coeruleus (Brochard et al., 2009; Cebrián et al., 2014; Gate et al., 2021; Sommer et al., 2018), neurons that are killed in PD. We have demonstrated T cell autoreactivity towards α-synuclein peptides in PD patients (Sulzer et al., 2017) and recently showed that peripheral α-synuclein-reactive T cells are most abundant in early stage PD patients (Lindestam Arlehamn et al., 2020). Further evidence suggests that T cell infiltration of the midbrain parenchyma is more common in early stage PD (Galiano-Landeira et al., 2020), but the precise role of T cells in PD pathophysiology remains unknown.

Interferon gamma (IFN-γ) is a proinflammatory cytokine released primarily from CD4+ T helper type 1 (Th1) CD4+ T cells, CD8+ T cells, and natural killer (NK) cells (Schroder et al., 2004), cell types with low abundance in the brain parenchyma. IFN-γ elicits cellular responses by increasing transcription of interferon-stimulated genes (ISGs), many of which encode transcription factors, antigen processing and presentation machinery, and innate immune effector proteins (Schroder et al., 2004). Although IFN-γ was formerly considered to be generally undetectable in the CNS (Popko et al., 1997), low levels of IFN-γ derived from meningeal T cells can regulate neuronal activity and behavior in healthy mice (Filiano et al., 2016). CNS cells may be exposed to high levels of IFN-γ under infectious and inflammatory conditions that feature blood brain barrier disruption and T cell infiltration (Popko et al., 1997). In a mouse model of viral encephalitis, the release of IFN-γ from CD8+ T cells has been suggested to drive the stripping of neuronal synapses via actions of phagocytic cells (Di Liberto et al., 2018; Kreutzfeldt et al., 2013).

Several reports indicate that IFN-γ may play roles in PD pathogenesis. Elevated plasma levels of IFN-γ were found in PD patients (Mount et al., 2007), and we recently identified altered transcriptional signatures in peripheral T cells of PD patients (Dhanwani et al., 2022) that may contribute to Th1/IFN-γ bias (Kustrimovic et al., 2018). Elevated levels of IFN-γ were reported in the substantia nigra, caudate, and putamen in PD, regions that contain dopaminergic cell bodies and their axonal projections (Mogi et al., 2007). Constitutive expression of IFN-γ in the lateral ventricles of mice leads to progressive nigrostriatal degeneration (Chakrabarty et al., 2011). Collectively, these data suggest that infiltrating T cells may contribute to dopaminergic demise in PD (reviewed in Garretti et al., 2022), in part by IFN-γ-driven neuroinflammation (Barcia et al., 2011; Ferrari et al., 2021).

Key questions on the possible role of IFN-γ in PD pathogenesis include: how do specific CNS cell types respond to IFN-γ, and which are capable of antigen presentation? Do midbrain neurons directly respond to IFN-γ (Cebrián et al., 2014) or are they bystanders to glial reactions (Mount et al., 2007)? In this study, we address these questions by analyzing neuronal and glial gene expression responses to IFN-γ injected into the ventral midbrain of adult mice. We find that MHC-I antigen processing and presentation genes are rapidly induced in all CNS cell types, including all major groups of midbrain neurons. Single nucleus RNA-sequencing (snRNA-seq) revealed numerous transcriptional responses that were restricted to glial cells, as well as microglia-specific expression of MHC-II antigen presentation genes. We show that dopamine neuronal responses to IFN-γ are abolished by the selective deletion of the IFN-γ-binding domain of IFNGR1, indicating that neurons are capable of MHC-I upregulation via cell-autonomous innate immune signaling.

## Results

### Kinetics of transcriptional responses to IFN-γ by midbrain neurons and glia

We chose to study the effects of intraparenchymal injection of IFN-γ to exclude effects of systemic inflammation or activation of the peripheral immune system, as previously reported (Gottfried-Blackmore et al., 2009). We first sought to measure the kinetics of IFN-γ responses in ventral midbrain neurons and glia. We delivered either 400 nL of 0.9% saline or 0.9% saline + 100 ng/μL IFN-γ (40 ng of IFN-γ) to the ventral midbrain bilaterally via stereotactic injection and sacrificed mice 6, 24, 48, and 72 hours after injection (**Fig. 1a**). After rapid tissue dissection, we prepared nuclei, conducted fluorescence-activated nuclear sorting (FANs) to separate neuronal nuclei (NeuN+) from other cell types (NeuN-), and analyzed the sorted nuclei by bulk RNA sequencing (RNA-Seq, **Supp. Fig. 1a**). Comparison of NeuN- and NeuN+ nuclei demonstrated substantial enrichment and depletion of pan-neuronal markers (e.g., *Syt1, Snap25, Rbfox3*), genes specific to excitatory, inhibitory, and dopaminergic neurons (e.g., *Slc17a6, Gad2, Slc18a2*), and genes specific to oligodendrocytes, astrocytes, and microglia (**Supp. Fig. 1b-c**). Hereafter, we refer to the NeuN-fraction as ‘glia’, as the prevalence of endothelial and mural cells within this fraction is exceedingly low compared to glial cell types (see **Fig. 3** below). Principal component analysis (PCA) revealed clear segregation of NeuN- and NeuN+ populations along PC1, which captured 80% of the variance (**Fig. 1b**). PC2 captured 5% of the variance and separated nuclei according to time of IFN-γ exposure, with the greatest differences observed at 6 and 24 hours for both neurons and glia (**Fig. 1b**). These results demonstrate that IFN-γ response kinetics are highly reproducible across replicates.

**Figure 1:**
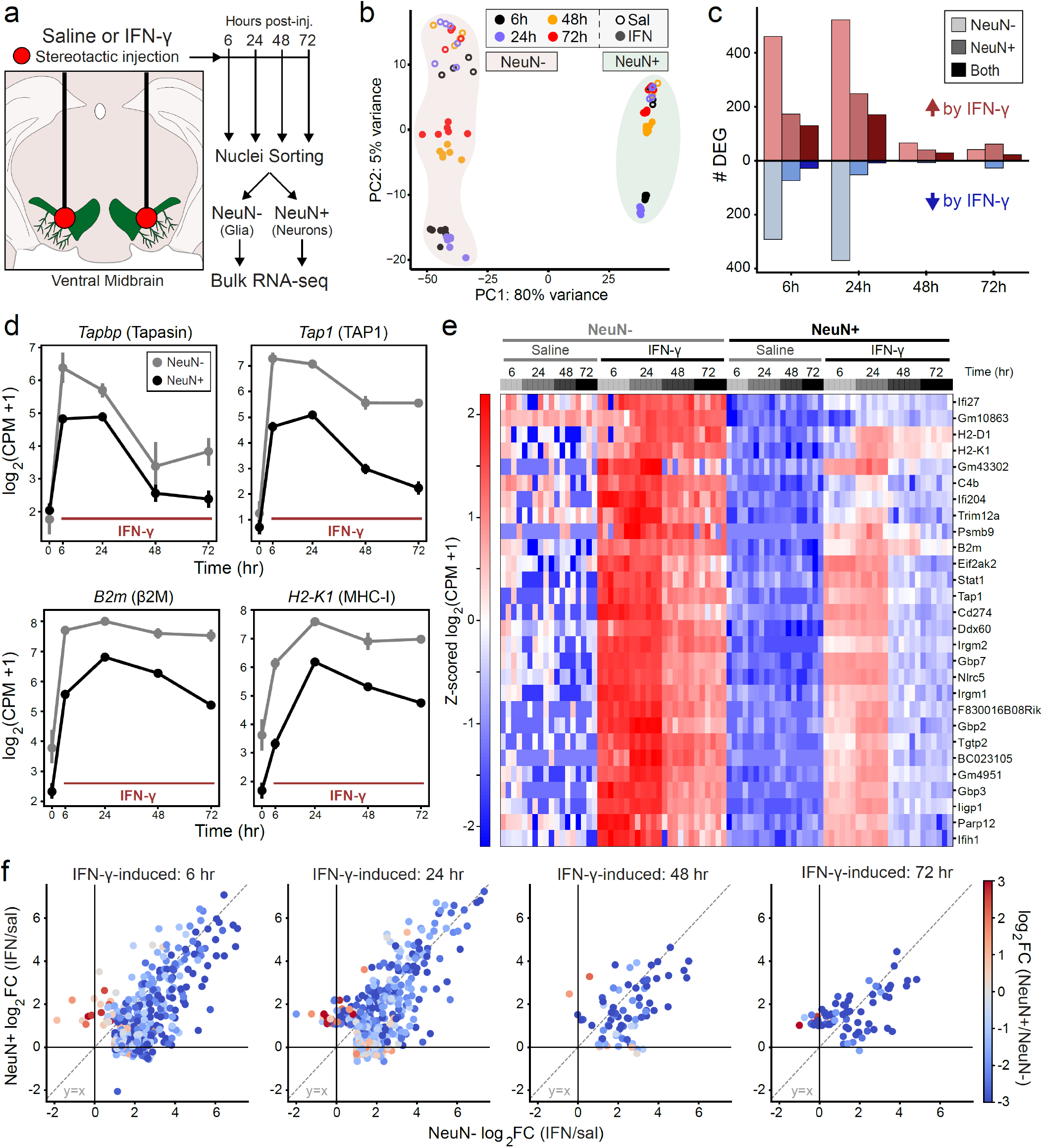
Core transcriptional responses to IFN-γ are conserved in neurons but are lower in magnitude compared to glia. **(a)** Experimental schematic for NeuN-/NeuN+ RNA-seq after injection of saline or IFN-γ. **(b)** Principal component analysis (PCA) of RNA-seq UMI counts normalized with *DESeq2 variance stabilizing transformation*. Each point represents a bulk RNA-seq replicate of ∼1000 sorted nuclei (n = 4-6 replicates each of NeuN-or NeuN+ nuclei from 2-3 mice for each treatment/timepoint). **(c)** Number of differentially expressed genes (DEG) for IFN-γ vs. Sal comparisons in NeuN-or NeuN+ samples, or both (overlap), at each timepoint (*DESeq2*, |log2FC| > 1 and pAdj < 0.01). **(d)** Mean ± SEM for mRNA abundance in NeuN-/NeuN+ samples, normalized as log2(CPM + 1). Samples as in (b), (n = 4-6 replicates each of NeuN-/+ nuclei from 2-3 mice for each treatment/timepoint). Saline samples from all timepoints are together at t=0. **(e)** Heatmap of z-scored, normalized mRNA abundance (log2[CPM + 1]) for 28 mRNAs upregulated at >3 timepoints in both NeuN- and NeuN+ samples (see **Supp. Fig. 1d**). Samples as in (b); each column is a replicate. **(f)** Scatter plots comparing the effect of IFN-γ (log2 fold change) in NeuN+ vs. NeuN-samples for all IFN-γ-induced genes at each timepoint (*DESeq2*, log2FC > 1 and pAdj < 0.01). Points are colored by differential expression in IFN-γ-treated NeuN+ vs. NeuN-samples (*DESeq2* log2FC).

Binding of IFN-γ dimers to the IFN-γ receptor (IFNGR) leads to cytoplasmic activation of Janus kinases 1-2, a cascade of tyrosine phosphorylation events, recruitment and phosphorylation of STAT1α, and nuclear translocation of phospho-STAT1α dimers (pSTAT1, reviewed in Schroder et al., 2004). pSTAT1 binding at γ-activation site (GAS) elements leads to increased transcription of ISGs involved in antiviral and antibacterial immunity, including other transcription factors. The first wave of rapidly-induced genes includes

IFN regulatory factor 1 (IRF-1), a transcription factor and master regulator of IFN-γ-induced gene expression (Namiki et al., 2005) that drives subsequent waves of transcription via binding to IFN-stimulated response elements (ISRE) (Schroder et al., 2004). Comparing IFN-γ to saline at each timepoint, we found the highest number of differentially expressed genes (DEGs; *DESeq2*, |log_2_FC| > 1, p_adj_ < 0.01) at 6 and 24 hours for both neurons and glia (**Fig. 1c**). At 6 and 24 hours, there were more than twice as many upregulated genes in glia compared to neurons, with considerable overlap in the upregulated genes (**Fig. 1c**). The number of DEGs in neurons and glia at 48 and 72 hours was comparatively modest, consistent with the decay of IFN-γ and/or negative feedback.

Consistent with prior kinetic analyses (Min et al., 1996), the induction of the ER peptide transporter mRNAs *Tap1* and *Tapbp* was rapid, with peak expression at 6 hours in both neurons and glia (**Fig. 1d**). MHC-I heavy chain (*H2-K1, H2-D1*) and light chain (*B2m*) genes do not harbor GAS elements, but rather are induced by IRF-1 binding to ISRE sequences in later waves of transcription (van den Elsen et al., 2004; Zhou, 2009). Accordingly, levels of *H2-K1* and β2m mRNAs were highest at 24 hours and remained ∼8-fold elevated at 72 hours (**Fig. 1d**). The rise and decay of all four mRNAs was markedly similar across neuronal and glial nuclei, although the amplitude of induction was higher in glia (∼2-8-fold higher expression relative to neurons at all timepoints, **Fig. 1d**).

Of the DEGs upregulated by IFN-γ in both neuronal and glial nuclei, 28 genes were upregulated at >3 timepoints (**Supp. Fig. 1d**), indicating sustained upregulation across multiple cell types and central involvement in the IFN-γ host defense program. Indeed, gene ontology (GO) analysis of these 28 genes identified top over-represented ontologies including ‘*Cellular response to IFN-γ*’, ‘*MHC class I protein complex*’, and ‘*Antigen processing and presentation of exogenous peptide antigen via MHC class I, TAP-dependent*’ (**Supp. Fig. 1d**). The induced genes included canonical IFN-inducible proteins such as MHC class I (*H2-K1, H2-D1, B2m*), immunoproteasome subunits (*Psmb9*/LMP2), ER peptide transporters (*Tap1*), guanylate binding proteins, and key regulators of IFN-γ-induced transcription (*Stat1, Nlrc5*) (**Fig. 1e**). Expression of these core IFN-γ-responsive genes was virtually absent in neuronal nuclei from saline-treated mice, and relative expression was uniformly higher in glia than in neurons (**Fig. 1e**). This phenomenon was true for most genes induced by IFN-γ at all time points: while the magnitude of upregulation by IFN-γ is well-correlated between neurons and glia, expression in glia was typically 2-8-fold higher than in neurons (**Fig. 1f**). These results demonstrate that the canonical IFN-γ transcriptional response occurs in both neurons and glia with similar kinetics, but with lower amplitude in neurons.

### IFN-γ induces expression of T cell chemoattractant and CCR2 ligand chemokines

Given the role of chemokines in regulating leukocyte migration into the CNS (Babcock et al., 2003), we analyzed the expression of mRNAs encoding chemokines. Several chemokines were expressed primarily in neurons and were unchanged after treatment with IFN-γ, including *Cx3cl1*/Fractalkine, *Ccl25/*TECK, and *Ccl27a/*CTACK (**Supp. Fig. 2a-b**). Neuronally derived CX3CL1/Fractalkine is known to modulate microglia via CX3CR1 (Harrison et al., 1998), while CCL25/TECK and CCL27/CTACK are primarily known for T cell homing in the gut and skin (Morales et al., 1999; Svensson et al., 2002) and to our knowledge have not been described in the CNS. In addition to well-known T cell chemoattractants (*Cxcl9*/MIG, *Cxcl10*/IP-10, *Cxcl11*/I-TAC), IFN-γ also induced the expression of CCR2 ligands (*Ccl2*/MCP1, *Ccl12*/MCP-5) in both neurons and glia, with levels peaking at 6 hours and declining rapidly thereafter (**Supp. Fig. 2c**). Consistent with prior studies (Di Liberto et al., 2018; Howe et al., 2017), these results suggest that IFN-γ-exposed ventral midbrain neurons express chemokines that attract T cells and modulate phagocytic function of microglia/macrophages.

### Cell type-specific temporal responses to IFN-γ in midbrain neurons and glia

We next used a generalized linear model in *DESeq2* to identify genes that are differentially regulated by IFN-γ in neurons vs. glia (NeuN vs. Drug interaction, **Supplementary Table 1**, see **Methods**). Forty-two genes were upregulated in both neurons and glia, but the effect of IFN-γ was greater in glia; these included canonical IFN-inducible genes such as *Tap2* and *Gbp2* (**Fig. 2a-c**, purple module). Most of the remaining DEGs from the interaction analysis were largely unchanged in neurons despite upregulation or downregulation by IFN-γ in glia (**Fig. 2a**). These genes could be further separated into three groups based on the effect of IFN-γ in glia and differences in baseline (saline) expression in neurons and glia.

**Figure 2:**
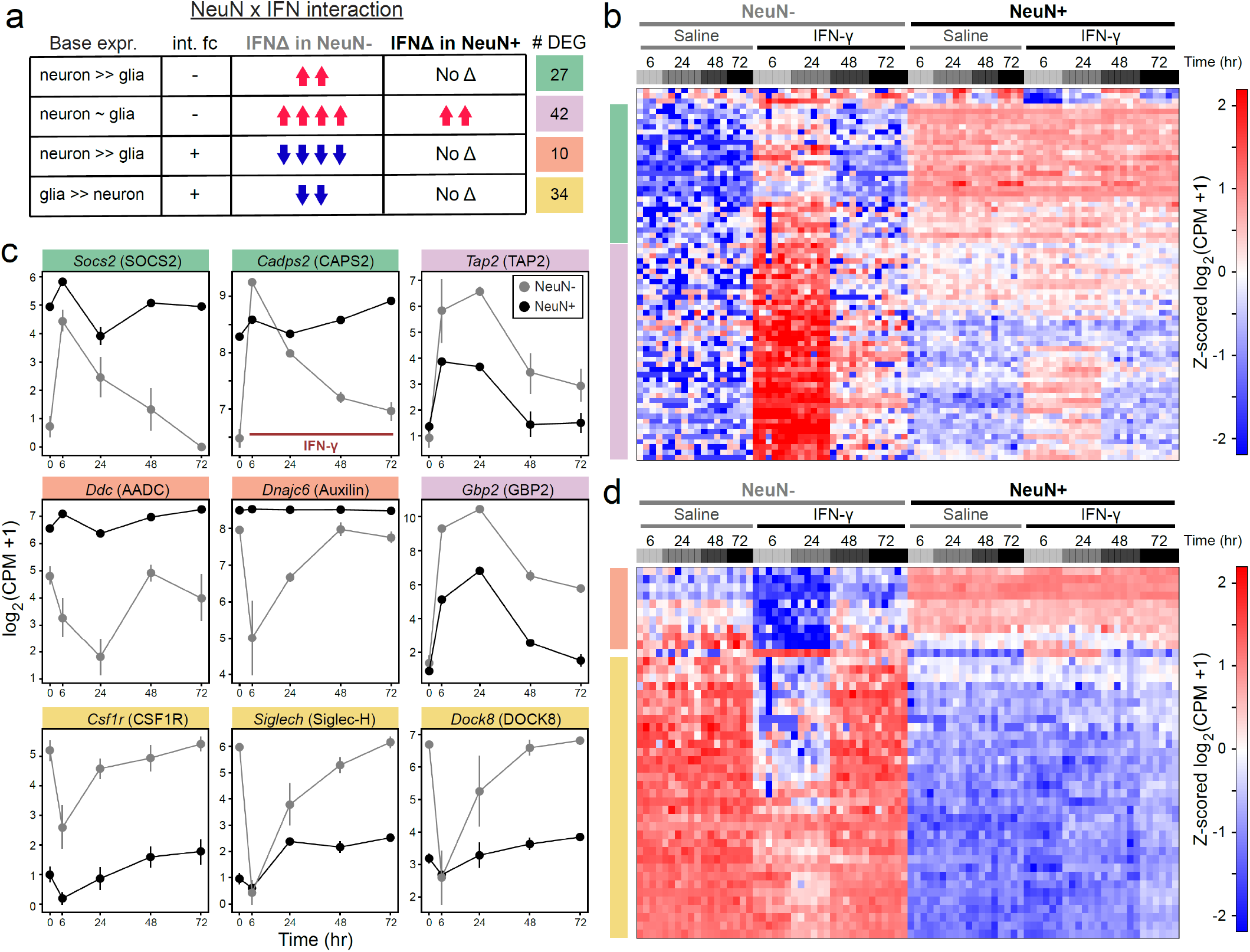
Divergent transcriptional responses to IFN-γ in glia. **(a)** Summary of DEGs from interaction analysis (*DESeq2*: ∼NeuN + IFN + NeuN:IFN), broken down by baseline expression and effect of IFN-γ in NeuN+ and NeuN-samples. **(b)** Heatmap of z-scored mRNA abundances for genes upregulated in NeuN-samples and significant in NeuN x IFN interaction analysis in (a), normalized as log2(CPM + 1). Each column represents a bulk RNA-seq replicate of ∼1000 sorted nuclei (n = 4-6 replicates from 2-3 mice for each treatment/timepoint). **(c)** Mean ± SEM for mRNA abundance in NeuN-/NeuN+ samples, normalized as log2(CPM + 1). Samples as in (b), (n = 4-6 replicates each of NeuN-/+ nuclei from 2-3 mice for each treatment/timepoint). All saline samples are together at t=0. **(d)** Heatmap of z-scored mRNA abundances for genes downregulated in NeuN-samples and significant in NeuN x IFN interaction analysis in (a), normalized as log2(CPM + 1). Each column represents a bulk RNA-seq replicate of ∼1000 sorted nuclei (n = 4-6 replicates from 2-3 mice for each treatment/timepoint).

Twenty-seven genes were highly expressed in neurons at baseline, but not glia, and were upregulated by IFN-γ only in glia (**Fig. 2a-c**, green module). These included the vesicular secretion regulator *Cadps2* (CAPS2) (Sadakata et al., 2004) and Socs2 (SOCS2), a negative feedback regulator of JAK-STAT signaling (Rico-Bautista et al., 2006) (**Fig. 2c**). Another 10 genes were also highly expressed in neurons at baseline, but not glia, but were rapidly downregulated by IFN-γ only in glia; these included the enzyme dopa decarboxylase (Ddc) that is required for dopamine synthesis, and the endocytic modulator Auxilin (*Dnajc6*) which is associated with rare cases of familial PD (**Fig. 2a & 2c-d**, orange module). These results suggest that glia, but not neurons, transiently suppress transcription of specific mRNAs involved in vesicular secretion and recycling after exposure to IFN-γ.

Finally, 34 genes were highly expressed in glia at baseline, but not neurons, and were rapidly downregulated by IFN-γ (**Fig. 2a & 2c-d**, yellow module). Notably, these included many microglia-enriched genes such as *Csf1r, Siglech*, and *Dock8* (**Fig. 2c**). We further found that 23 of the top 50 microglial marker genes (identified via snRNA-seq data, see **Figure 3** below) were significantly downregulated at 6 hours after IFN-γ (**Supp. Fig. 3**). Thus, the microglial activation in response to IFN-γ likely involves a transient repression of some, but not all, lineage marker genes.

**Figure 3:**
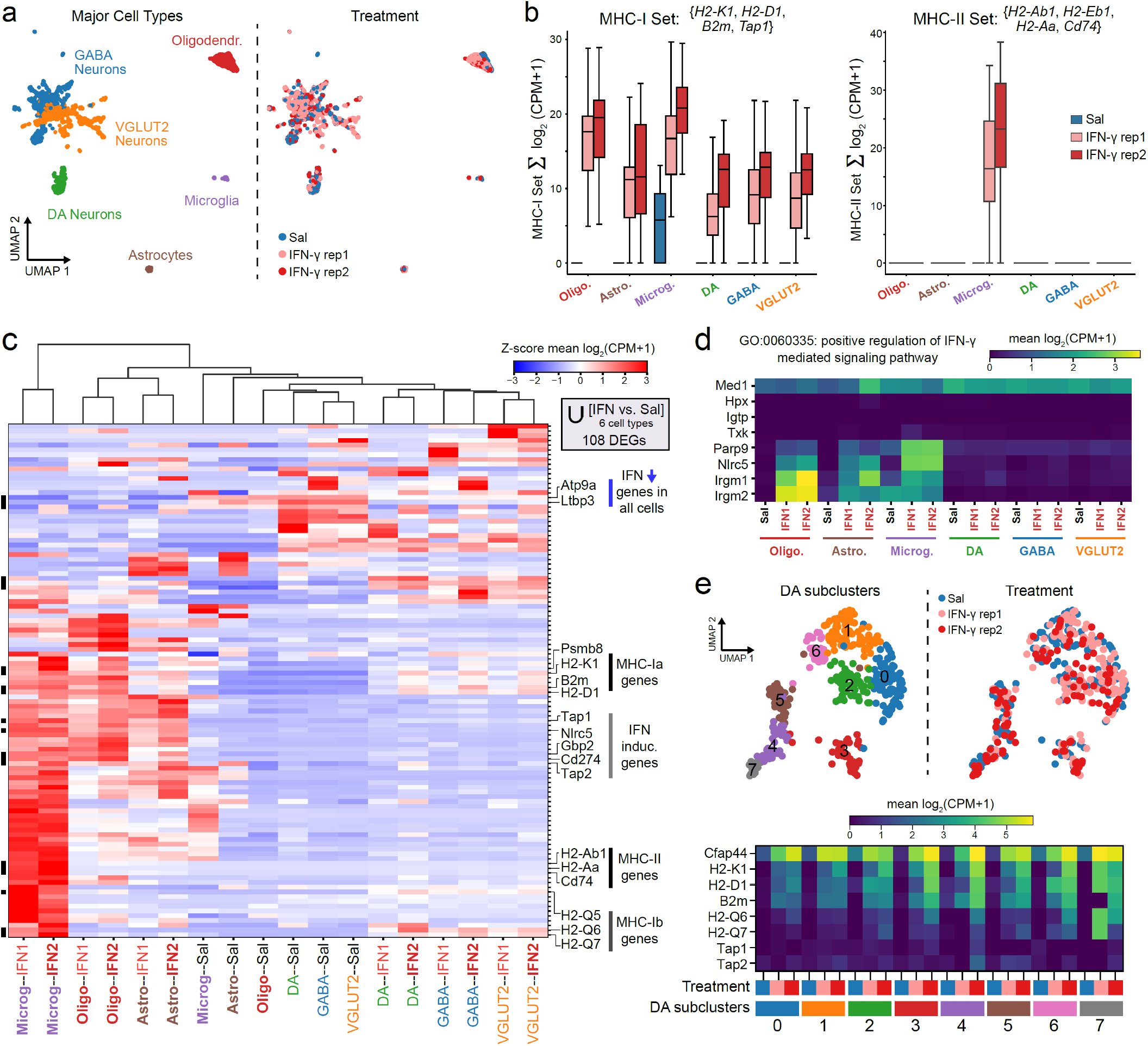
Single nucleus RNA-seq analysis of responses to IFN-γ. **(a)** UMAP embedding of snRNA-seq profiles from the ventral midbrain used for downstream analysis (4,699 nuclei after collapse to major cell types and filtering, see **Methods**). *Left*: major cell types for downstream analysis, see **Supp. Fig. 4b** for clustering with marker genes. *Right*: mouse sample origin (n = 1 for saline, n = 2 for IFN-γ). **(b)** Box and whiskers plots depicting the sum of mRNA abundance (log2[CPM+1]) in each major cell group for the indicated sets of MHC-I or MHC-II genes. **(c)** Clustered heatmap of z-scored mRNA abundance (log2[CPM+1]) for the union of highly differentially expressed genes across all major cell groups (modified Mann-Whitney U-test comparing IFN vs. Sal, |log2FC| > 4 and q < 0.05). **(d)** Heatmap of average mRNA abundance (log2[CPM+1]) for genes in MGI GO:0060335: ‘positive regulation of interferon-gamma-mediated signaling pathway’. **(e)** *Upper*: UMAP embedding of dopamine neuronal snRNA-seq profiles with subcluster IDs and mouse sample origin. *Lower*: Heatmap of average mRNA abundance (log2[CPM+1]) for select IFN response genes within dopamine neuronal subclusters from each mouse sample.

### Conserved and cell-type-specific single cell responses to IFN-γ

To resolve IFN-γ responses at the single cell level, we next conducted snRNA-seq on ventral midbrain nuclei. To ensure proper coverage of neuronal subtypes, we used FANS as above to enrich NeuN+ nuclei to ∼70% of each mouse sample (n=1 saline, n=2 IFN-γ) prior to sequencing. Since the core IFN-γ-induced signature was stably upregulated for at least 48 hours (**Fig. 1**), we selected 48 hours for subsequent analysis in order to better approximate chronic *in vivo* exposure and avoid transient disruption of cell lineage marker expression (**Fig. 2c-d**). We processed the sequencing data and performed unsupervised clustering on the molecular count matrices as described previously (Levitin et al., 2019; Yuan et al., 2018). We identified thirty-eight clusters (**Supp. Fig. 4a**), which were collapsed to six major cell types with sufficient coverage for downstream analysis based on key marker genes significantly enriched in each cluster (**Supp. Fig. 4b**). After removal of nuclear doublets (**Supp. Fig. 4c-d**, see **Methods**), we retained a total of 4,699 nuclei, including 1,241 oligodendrocytes, 194 astrocytes, 134 microglia, 437 dopamine neurons, 1,257 GABAergic neurons, and 1,436 VGLUT2 neurons for downstream analysis (**Fig. 3a**), with comparable distribution across saline- and IFN-γ-treated mice (**Supp. Fig. 5a**).

We previously found that IFN-γ-treated dopamine neurons express MHC-I *in vitro* (Cebrián et al., 2014), but dopamine neurons comprise a minority of all ventral midbrain neurons represented in our bulk NeuN+ RNA- seq data (**Fig. 1**). We first analyzed the expression of a gene signature of MHC-I related genes (*H2-K1, H2-D1, B2m, Tap1*). In the absence of IFN-γ, only microglia expressed detectable levels of these genes, whereas all cell types could be induced to express the MHC-I gene signature (**Fig. 3b**). We previously reported that non-catecholaminergic neurons express MHC-I with higher concentrations of IFN-γ *in vitro* (Cebrián et al., 2014), and consistently, we found that GABAergic, glutamatergic, and dopaminergic midbrain neurons all expressed similar levels of MHC-I genes after exposure to high dose IFN-γ *in vivo* (**Fig. 3b**). Induced levels of MHC-I genes were similar between astrocytes and neurons, all of which were lower than in oligodendrocytes and microglia (**Fig. 3b**). Thus, all major cell types in the mouse ventral midbrain parenchyma can upregulate MHC-I mRNA upon exposure to IFN-γ.

In contrast to MHC-I, the expression of MHC-II is typically limited to professional antigen presenting cells (van den Elsen et al., 2004). We found that the MHC-II genes (*H2-Ab1, H2-Eb1, H2-Aa*) and the invariant chain (Cd74) were highly expressed in IFN-γ-treated microglia, and no other cell types showed detectable MHC-II gene expression (**Fig 3b, Supp. Fig. 5c**). Similarly, we found that IFN-γ led to marked changes in the morphology of Iba1^+^ microglia, with associated cellular proliferation as detected by Ki67 protein expression and nuclear localization (**Supp. Fig. 6**). Ki67 staining was not present within other cell types.

We next conducted differential expression analysis, comparing each IFN-γ vs. saline sample within each major cell type. Only genes differentially regulated by IFN-γ in both samples were retained (see **Methods**; |log_2_FC| > 3, FDR < 0.01), which comprised 108 DEGs across all six major cell types (**Fig. 3c**). Hierarchical clustering revealed that the most distinct expression profile for these genes belonged to IFN-γ-activated microglia, followed by oligodendrocytes and astrocytes, while the profiles of saline- and IFN-γ -exposed neurons were more similar (**Fig. 3c**). We found that two genes, *Atp9a* and *Ltbp3*, were massively downregulated in all cell types (**Supp. Fig. 5b**). ATP9A is a type IV P-type ATPase involved in endosome recycling and exosome release (Naik et al., 2019; Tanaka et al., 2016), while LTBP-3 is a regulator of TGF-β (Robertson et al., 2015), although to our knowledge neither has been reported to possess an explicit immune function. Meanwhile, the majority of genes were upregulated by IFN-γ and were highly expressed in glia, but not neurons (**Fig. 3c**). One notable exception to this trend was the expression of the non-classical MHC-I (class Ib) genes *H2-Q5, H2-Q6*, and *H2-Q7*, which were robustly induced by IFN-γ in microglia and all neurons, but not oligodendrocytes and astrocytes (**Fig. 3c**). Class Ib molecules are structurally similar to class Ia (*H2-D1, H2-K1*), although class Ib genes tend to exhibit tissue-specific expression and distinct regulatory elements (Howcroft and Singer, 2003).

While all neurons upregulated classical MHC-I genes, we found that a number of canonical ISGs (e.g., *Tap1, Tap2, Gbp2, Cd274*/PD-L1) were poorly expressed in IFN-γ-treated neuronal nuclei at 48 hours (**Fig. 3c**). Notable among these was *Nlrc5*, which encodes NLRC5, a member of the NOD-like receptor family that critically regulates MHC-I expression analogous to CIITA regulation of MHC-II expression (Jongsma et al., 2019). These results suggest that deficiency in effectors of IFN-γ-mediated signaling or transcription might lead to the dampened neuronal responses observed in our studies. Indeed, we found that several genes associated with the GO term ‘positive regulation of IFN-γ mediated signaling pathway’ were robustly induced in glia, but not in neurons (**Fig. 3d**). These included poly(ADP-ribose) polymerase 9 (PARP9), which is known to activate STAT1 phosphorylation in activated macrophages (Iwata et al., 2016), and immunity-related GTPases *Irgm1-2* which regulate IFN-γ-driven effector responses to intracellular pathogens (Taylor, 2007).

Within the ventral midbrain of PD patients, dopamine neurons of the ventral tegmental area (VTA) tend to be spared relative to more vulnerable populations in the SNc (Hirsch et al., 1988). Notably, we previously found HLA class I expression in dopamine neurons within the SNc, but not VTA, in postmortem tissue from PD patients (Cebrián et al., 2014). However, recent studies have demonstrated considerable transcriptomic heterogeneity amongst midbrain dopamine neurons (Poulin et al., 2014; Saunders et al., 2018). We thus leveraged our snRNA-seq data to study transcriptional responses to IFN-γ amongst various subpopulations of dopamine neurons. We further analyzed dopamine neuronal expression profiles using unsupervised clustering as above (see **Methods**) and identified eight subclusters (**Fig. 3e** and **Supp. Fig. 7**). All subclusters expressed moderate to high levels of *Slc6a3* (DAT) but were distinguished by expression of marker genes consistent with previous studies (*Ndnf, Ntng1, Nefl, C1ql3, Cbln1, Cbln4, Tacr3*), including those with ventral SN bias (*Sox6, Aldh1a1, Grin2c*, and *Atp2a3*) and those with medial VTA bias (*Sorcs3, Nrp2, Grp, Calb1*) (Chung et al., 2005; Hobson et al., 2022b; Poulin et al., 2014; Saunders et al., 2018) (**Fig. 3e, Supp. Fig. 7**, and **Supplementary Table 2**). Dopamine neurons from saline- and IFN-γ-treated mice were distributed similarly throughout all subclusters (**Fig. 3e**), suggesting that IFN-γ treatment did not significantly alter the expression of subcluster-defining marker genes. We found no evidence of differential responses to IFN-γ, with all subclusters upregulating classical (*H2-K1, H2-D1, B2m*) and non-classical MHC-I genes (*H2-Q6, H2-Q7*) (**Fig. 3e**). Thus, at least under conditions of direct exposure to high concentrations of IFN-γ, an MHC-I antigen presentation program can be induced in all populations of ventral midbrain dopamine neurons.

### Dopamine neurons respond to IFN-γ via cell autonomous IFNGR signaling

In addition to IFN-γ, MHC-I heavy chain transcription can be induced in neurons by TNF-α (Neumann et al., 1997), and it is possible that the effects of IFN-γ on neurons may be due to release of other cytokines or interactions with glia, particularly activated microglia (Mount et al., 2007). Furthermore, canonical ISGs can be upregulated in dopamine neurons without IFN-γ under conditions of oxidative stress, such as high dose L-DOPA (Cebrián et al., 2014), 6-OHDA exposure (Mo et al., 2018), or a model of α-synuclein aggregation (Ugras et al., 2018). Thus, we first confirmed that dopamine neurons in the ventral tegmental area (VTA) and SNc express IFN-γ receptor subunit 1 (*Ifngr1*) mRNA using fluorescence in situ hybridization (FISH) (**Supp. Fig. 8**). To test whether cell autonomous IFN-γ signaling is required for their transcriptional responses, we selectively modified the *Ifngr1* locus in dopamine neurons by crossing *Ifngr1*^fl/fl^ mice to dopamine transporter (DAT)-cre mice. As previously described, Cre-mediated recombination in *Ifngr1*^fl/fl^ mice leads to deletion of the IFN-γ-binding extracellular domain of IFNGR1 and conditional loss of IFN-γ signaling (Lee et al., 2013).

We first confirmed that DAT-Cre:*Ifngr1*^fl/fl^ mice showed no gross alteration in midbrain or striatal TH and DAT immunolabel (**Supp. Fig. 9**). We then used immunohistochemistry to analyze phosphorylation of STAT1 Tyr701 (pSTAT1-Y701), the STAT1 residue critical for IFN-γ-mediated nuclear translocation and transcriptional activation (Shuai et al., 1993), in the ventral midbrain at 6 hours after injection of IFN-γ or saline. IFN-γ-induced nuclear pSTAT1-Y701 was observed in TH^+^ dopamine neurons and TH^-^ cells in WT:*Ifngr1*^fl/fl^ mice, but only in TH^-^ cells in DAT-Cre:*Ifngr1*^fl/fl^ mice (**Fig. 4a**). Triple labeling with NeuN enabled analysis of nuclei from dopamine neurons (TH^+^/NeuN^+^), non-dopamine neurons (TH^-^/NeuN^+^), and glia (TH^-^/NeuN^-^) (**Fig. 4b**). Quantification of pSTAT1-Y701 induction in these populations confirmed a selective deficiency of IFN-γ response in dopamine neurons (**Fig. 4c**). Notably, the average pSTAT1-Y701 intensity in glia was ∼3-fold higher than in neurons, with no appreciable differences between dopamine neurons and non-dopamine neurons (**Fig. 4b-c**). Consistent with the pSTAT1-Y701 results, RNA-FISH confirmed that dopamine neurons in IFN-γ-treated DAT-Cre:*Ifngr1*^fl/fl^ mice express significantly less *Tap1* and MHC-I heavy chain mRNAs compared to WT:*Ifngr1*^fl/fl^ mice (**Fig. 4e-f**). Collectively, these results confirm that cell autonomous IFNGR signaling and nuclear pSTAT1 translocation, not bystander responses to reactive glia, are responsible for transcription of antigen presentation genes within dopamine neurons following ventral midbrain exposure to IFN-γ.

**Figure 4:**
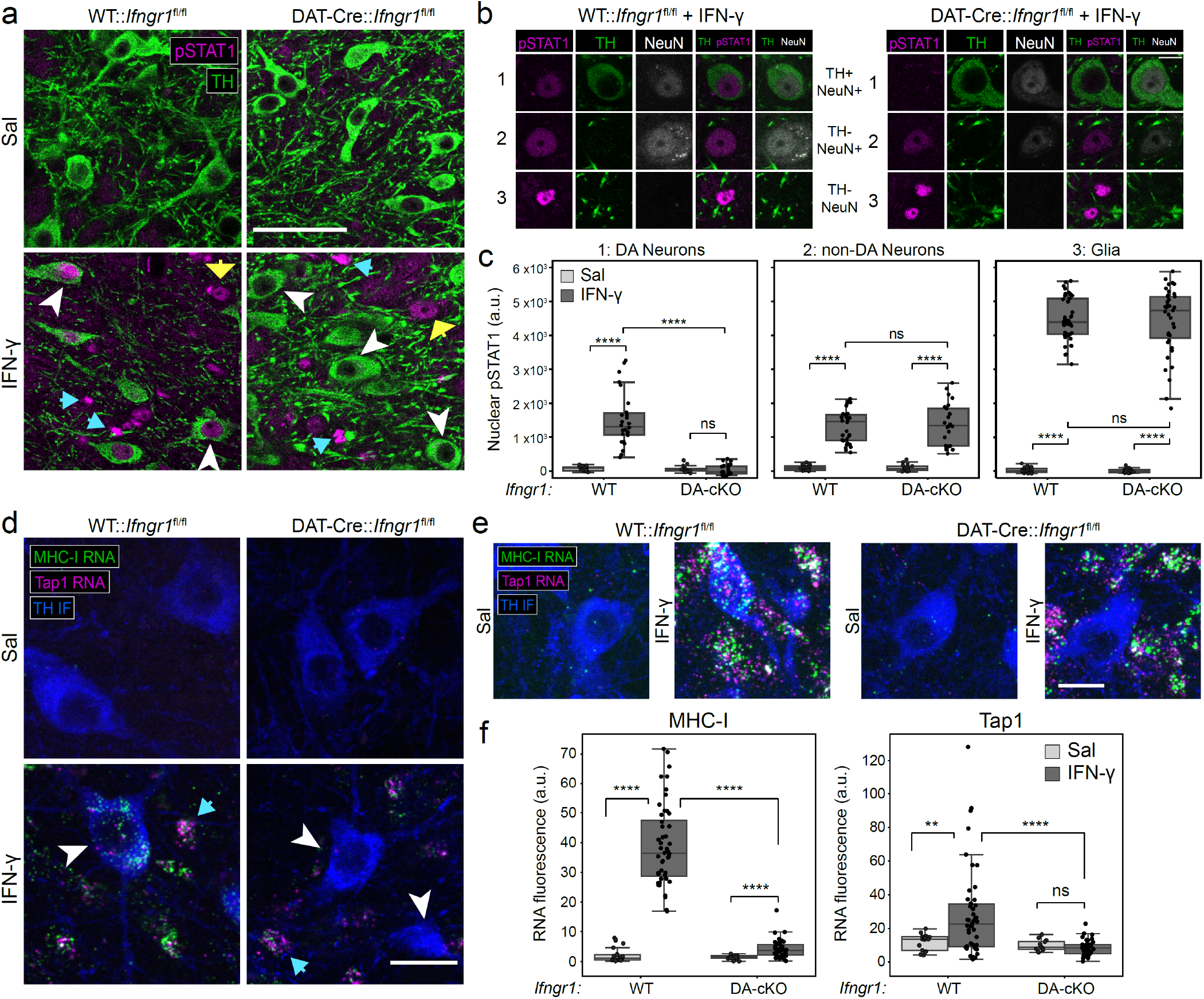
*Ifngr1* is required for pSTAT1 induction and MHC-I upregulation in mDA neurons after *in vivo* exposure to IFN-γ. **(a)** TH and pSTAT1-Y701 immunofluorescence in the SNc of WT:Ifngr1^fl/fl^ or DAT-Cre:Ifngr1^fl/fl^ mice after saline or IFN-γ. White arrowheads indicate TH^+^ mDA neurons, blue arrows indicate NeuN^-^ nuclei, and yellow arrows indicate TH^-^ neurons. Scale bar: 50 μm. **(b)** pSTAT1-Y701 in TH^+^/NeuN^+^ mDA neurons, TH^-^/NeuN^+^ non-DA neurons, and TH^-^/NeuN^-^ glial nuclei in the SNc of WT:Ifngr1^fl/fl^ or DAT-Cre:Ifngr1^fl/fl^ mice after exposure to IFN-γ. Scale bar: 10 μm. **c)** Average nuclear pSTAT1 intensity (arbitrary units, median background subtracted) in each of the indicated cell types, genotypes, and treatment groups, related to (a-b). 10-20 neurons were quantified per region per mouse; n=2 saline, n=3 IFN-γ. **** p < 0.001, Mann-Whitney U test. (**d-e**) MHC-I and Tap1 RNA FISH in the SNc of WT:Ifngr1^fl/fl^ or DAT-Cre:Ifngr1^fl/fl^ mice after saline or IFN-γ. White arrowheads indicate TH^+^ mDA neurons, blue arrows indicate other cells with intact Ifngr1. Scale bars (d): 20 μm, (e): 15 μm. (**f**) Average MHC-I or Tap1 RNA intensity (arbitrary units, median background subtracted) in TH^+^ mDA neurons of the indicated genotype and treatment groups, related to (d-e). 10-20 neurons were quantified per region per mouse. WT; n=3 Sal, n=4 IFN-γ. DAT-Cre:Ifngr1^fl/fl^ (DA-cKO); n=3 Sal, n=4 IFN-γ. **** p < 0.001, ** p < 0.01, Mann-Whitney U test.

## Discussion

Interferons were discovered as factors with antiviral activity (Isaacs et al., 1957) and research over the following decades identified a large set of interferon-regulated pathways, including about 10% of the human genome (Samarajiwa et al., 2009). Due to reports of CD8+ and Th1 CD4+ T cells infiltrating the brain in aging and neurological disease, we characterized transcriptional changes caused by acute local exposure to the proinflammatory cytokine IFN-γ in the ventral midbrain. To avoid the confounding influence of other inflammatory cytokines, we designed our current approach to provide a molecular analysis of acute cellular changes in the absence of peripheral inflammation.

### Conserved responses to IFN-γ in brain cells

Here, we focused on the brain parenchymal response to IFN-γ and note that rare cell types in or at the border the CNS, including dendritic cells, monocytes, natural killer cells, and border associated macrophages were either too uncommon for detection or lost during brain dissection due to meningeal removal: these cell types may nevertheless play important roles in adaptive immune responses in the brain. Although MHC-I is expressed by most nucleated cells in the periphery, our results confirm previous reports that MHC-I mRNA is expressed at negligible levels in the healthy rodent CNS (Vass and Lassmann, 1990), with the exception of microglia (**Fig. 3b**). However, we find that for all major classes of CNS cells, IFN-γ rapidly induces expression of MHC-I and other ISGs involved in antigen presentation, with similar kinetics to non-brain cells (Min et al., 1996; van den Elsen et al., 2004). Transcriptional kinetics were similar across CNS cell types, but ISG expression amplitude was highest in microglia, lower in astrocytes and oligodendrocytes, and lowest in neurons (**Fig. 3**). However, the abundance of neurons compared to microglia suggests that neuronal expression may still be important for the overall brain response to IFN-γ. For example, the IFN-γ-inducible T cell chemoattractants *Cxcl9, Cxcl10, and Cxcl11* were rapidly upregulated in both neurons and glia (**Supp. Fig. 2**). Although our data suggest that microglia induce these genes at >8-fold higher levels than neurons, neurons are typically >5-fold more abundant than microglia in the human brain (Santos et al., 2020). Thus, it is likely that neurons, astrocytes, oligodendrocytes, and microglia each contribute to conserved IFN-γ responses in the brain.

### Dampened neuronal responses to IFN-γ

Why are neurons less responsive to IFN-γ than glia? It has been suggested that the absence of MHC-I expression and lack of antigen presentation capacity may be protective for post-mitotic neurons (Rall, 1998). Our results underscore the longstanding view that MHC-I is not readily detectable in neurons, and highlight a growing appreciation that detection of physiologically relevant neuronal MHC-I requires highly sensitive techniques (McDole et al., 2010).

Multiple mechanisms could underlie the dampened transcriptional response to IFN-γ in neurons. First, we observed that the levels of nuclear pSTAT1 at 6 hours post-IFN-γ are significantly lower in neurons compared to glia (**Fig. 4**). Decreased abundance of IFNGR, STAT1, and/or negative feedback signaling that regulates STAT1-Y701 phosphorylation could all contribute to decreased nuclear pSTAT1 accumulation (Rose et al., 2007). Given the dramatic response in microglia compared to other glia, differential regulation of pSTAT1 is unlikely to be the sole mediator of cell type-specific IFN-γ responses in the CNS. Other minor differences in ISG induction between cell types suggest subtle differences in transcriptional dynamics downstream of initial pSTAT1 binding: for example, the upregulation of non-classical MHC-I genes *H2-Q5, H2-Q6*, and *H2-Q7* was specific to microglia and neurons (**Fig. 3c**). Thus, the overall lower amplitude of IFN-γ-induced transcription in neurons could be due in part to altered basal chromatin accessibility and/or inefficient expression of key transcription factors that drive downstream expression programs.

### Microglia-specific responses to IFN-γ

Unexpectedly, we found an early, transient downregulation of some, but not all, microglial lineage-identity genes in NeuN-nuclei at 6 hours post-exposure to IFN-γ (**Supp. Fig. 3**).

These include *Dock8*, a protein implicated in controlling microglial function in neurodegeneration (Gunsolly et al., 2010), the cytokine receptors *Csf1r* and *Csf3r*, and *Siglech*, a sialic acid-binding receptor that inhibits microglial activation (Klaus et al., 2021). These responses may lead to reduced microglial sensitivity to anti-inflammatory and homeostatic mediators, as has been observed in IFN-γ-polarized macrophages (Ivashkiv, 2018). However, further studies are required to characterize the role of this transient transcriptional suppression within the microglial activation program induced by IFN-γ.

Within the CNS, it is well-established that activated microglia express MHC-II (McGeer et al., 1988; Vass and Lassmann, 1990), but prior studies of MHC-II expression in mouse oligodendrocytes and astrocytes have yielded conflicting results (Falcão et al., 2018; Rostami et al., 2020; Stüve et al., 2002; Vass and Lassmann, 1990). In the context of MS, single cell RNA-seq in mouse models (Falcão et al., 2018; Kaya et al., 2022) and human brain tissue (Jäkel et al., 2019) identified subsets of mature oligodendrocytes that express IFN-inducible genes, including MHC-I and

MHC-II genes, although only MHC-I associated signals was observed in human neuropathology (Höftberger et al., 2004). Our results suggest that acute stimulation with IFN-γ elicits the transcriptional upregulation of MHC-II genes only in microglia (**Fig. 3b**), although more chronic exposure or the presence of other stimulatory factors may lead to expression in astrocytes and/or oligodendrocytes under certain conditions.

### Dopamine neurons directly respond to IFN-γ

Several studies indicate that neuronal MHC-I presentation could be a response to intrinsic neuronal stress, even in culture paradigms lacking cell types that release IFN-γ, TNF-α, or other candidate extrinsic triggers. In our hands, cultured postnatal DA neurons can be induced to present an intracellularly processed foreign antigen by MHC-I following high cytosolic oxidative stress caused by high levels of L-DOPA (Cebrián et al., 2014), in the absence of microglia, T cells, or extrinsic cytokines. The hypothesis that neuronal stress may be an independent means to drive antigen presentation is consistent with reports of increased MHC-I in mature neurons by kainic acid, which indices excitotoxicity (Corriveau et al., 1998), and increased TAP-1 and immunoproteasome messages following the toxin 6-hydroxy-dopamine (Mo et al., 2018) and by aggregated alpha-synuclein (Ugras et al., 2018). However, we found that dopamine neurons in DAT-Cre:*Ifngr1*^fl/fl^ mice did not significantly upregulate pSTAT1 or MHC-I in response to IFN-γ injection in the ventral midbrain (**Fig. 4**). These results suggest that the acute stress placed on dopaminergic neurons by 48 hours of surrounding glial activation is insufficient to upregulate pSTAT1 via alternate pathways, which may act at longer time scales.

Although IFN-γ has been implicated in dopaminergic neurodegeneration, studies in rodent and culture models of PD have focused on glial activation as the key driver of neuronal cell death (Barcia et al., 2011; Mount et al., 2007). IFNGR1 and STAT1 knockout mice are protected from pathological overexpression of IFN-γ in the mouse brain (Strickland et al., 2018), but it is unknown which cells require IFN-γ signaling to drive nigrostriatal degeneration in this model. Consistent with

## Materials and Methods

### Animals

Mice were housed on a 12-hour light/dark cycle with food and water available *ad libitum.* Adult male and female mice (6-12 months old) were used in all experiments. DAT^IRES-Cre^ mice (JAX #006660, RRID: IMSR_JAX:006660) previous studies of cortical and hippocampal neurons (Filiano et al., 2016; Flood et al., 2019), we found that midbrain dopaminergic neurons express *Ifngr1* and upregulate MHC-I genes via increased nuclear pSTAT1-Y701 after IFN-γ exposure (**Fig. 4**). Although we directly exposed dopaminergic neurons to IFN-γ in the ventral midbrain, previous work found that IFN-γ stimulation of axons can induce neuron MHC-I via retrograde signaling (Clarkson et al., 2018). Given that their massive axonal arbors comprise the majority of their surface area (Matsuda et al., 2009) and protein mass (Hobson et al., 2022a), dopamine neurons may also be induced to express MHC-I via inflammation within the caudate and putamen in PD (Mogi et al., 2007).

The role of IFN-γ signaling and MHC-I expression within dopamine neurons in PD remains unclear, but evidence from mouse models of autoimmune-mediated narcolepsy, paraneoplastic encephalitis, and neurotropic viral infection suggest several possibilities. Bernard-Valnet et al. (2016) and Yshii et al. (2016) demonstrated that neuronal expression of ectopic “neo-self-antigen” led to antigen-specific CD8+ T cell infiltration of the brain parenchyma and neuronal destruction. While Kreutzfeldt et al. (2013) found that viral antigen-specific CD8+ T cell-mediated neuronal destruction requires functional IFN-γ receptor signaling in the CNS, constitutive neuronal overexpression of the MHC-I molecule H-2D^b^ did not elicit pathology in the absence of CNS IFN-γ receptor signaling. These results raised the possibility that neuronal upregulation of MHC-I is not critical for IFN-γ-mediated neuropathology in this model (Kreutzfeldt et al., 2013), but our results suggest that loss of neuronal IFNGR1 would completely prevent upregulation of the machinery necessary to process and present antigen on the constitutively expressed H-2D^b^ molecules (e.g., *B2m, Tap1, Psmb8*). Nonetheless, subsequent studies found that IFN-γ-mediated synapse loss in virally infected neurons was driven by neuronal CCL2 expression and phagocyte-mediated synaptic removal (Di Liberto et al., 2018). Thus, in addition to potential autoimmune-mediated responses directed at dopamine neurons, future work should also investigate the synaptic and electrophysiological consequences of IFN-γ signaling in the ventral midbrain.

(Bäckman et al., 2006) and Ifngr1^fl^ mice (JAX #025394, RRID: IMSR_JAX:025394) (Lee et al., 2013) were obtained from Jackson Laboratories (Bar Harbor, ME). DAT^IRES-Cre^ mice derive from C57BL/6J background, while Ifngr1^fl^ mice derive from C57BL/6N background. All experiments were conducted according to NIH guidelines and approved by the Institutional Animal Care and Use Committees of Columbia University and the New York State Psychiatric Institute.

DAT^IRES-Cre^:Ifngr1^fl^ experimental litters were bred from a cross of DAT^IRES-Cre/wt^;Ifngr1^fl/fl^ x DAT^wt/wt^;Ifngr1^fl/fl^ mice such that Cre^+^ mice (DAT^IRES-Cre/wt^;Ifngr1^fl/fl^) could be compared to Cre^-^ littermates with no modification of the Ifngr1 locus (DAT^wt/wt^;Ifngr1^fl/fl^). To ensure proper allocation of genotypes to experimental treatments (stereotaxic injection of virus and/or IFN-γ), experimenters were not blinded to the genotype of mice during stereotaxic injection. However, a separate experimenter that was blind to the genotype and treatment condition conducted animal sacrifice and tissue dissection. All experimental procedures were conducted according to NIH guidelines and were approved by the Institutional Animal Care and Use Committees of Columbia University and the New York State Psychiatric Institute.

### Stereotaxic Injections

All surgical procedures were approved by the Institutional Animal Care and Use Committee and the Department of Comparative Medicine at New York State Psychiatric Institute. Mice were anesthetized with 4% isoflurane.

Animals were transferred onto a Kopf Stereotaxic apparatus and maintained under isoflurane anesthesia (1–2%). After hair removal and sterilization of the scalp using chlorhexidine and ethanol, a midline incision was made. Bregma and Lambda coordinates were determined, and minor adjustments in head position were made to match the DV coordinates. Saline or IFN-γ was injected at AP –3.2, ML −0.9, and DV −4.4. A small hole was drilled into the skull and 400 nL of 0.9% saline or 0.9% saline + 100 ng/μL recombinant mouse IFN-γ (40 ng IFN-γ total) was injected through a pulled glass pipet using a Nanoject 2000 (Drummond Scientific; 8 pulses of 50 nL). At 5 minutes after injection, the glass pipet was slowly withdrawn over 5 min. After closing the skin with vicryl sutures, mice received 0.5 mL of 0.9% saline i.p. and were allowed to recover for >1 hour before being returned to their home cages.

### Antibodies and Reagents

| Name | Manufacturer | Catalog | RRID | Usage |
| --- | --- | --- | --- | --- |
| Chicken anti-TH | Millipore | AB9702 | RRID:AB_570923 | IHC, 1:500 |
| Mouse anti-NeuN | Millipore | MAB377 | RRID:AB_2298772 | IHC, 1:500 |
| Rabbit anti-Iba1 | Wako | 019-19741 | RRID:AB_839504 | IHC, 1:1000 |
| Rat anti-Ki67 | ThermoFisher | 14-5698-82 | RRID:AB_10854564 | IHC, 1:1000 |
| Mouse anti-NeuN, Alexa488 | Millipore | MAB377X | RRID:AB_2149209 | FANS, 1:500 |
| Rabbit anti-pStat-Y701 | Cell Signaling | 9167 | RRID:AB_561284 | IHC, 1:1000 |
| Rat anti-DAT | Millipore | MAB369 | RRID:AB_2190413 | IHC, 1:500 |
| Goat anti-Chicken IgY (H+L), Alexa Fluor Plus 488 | ThermoFisher | A-32931TR | RRID:AB_2866499 | IHC, 1:1000 |
| Goat anti-Mouse IgG (H+L), Alexa Fluor Plus 568 | ThermoFisher | A-11004 | RRID:AB_2534072 | IHC, 1:1000 |
| Goat anti-Rabbit IgG (H+L), Alexa Fluor 647 | ThermoFisher | A32733 | RRID:AB_2633282 | IHC, 1:1000 |
| Goat anti-Rat IgG (H+L), Alexa Fluor 647 | ThermoFisher | A-32931TR | RRID:AB_2866499 | IHC, 1:1000 |
| Mm-*Ifngr1*-C2 | ACD Biotechne | 441491-C2 | NA | FISH |
| Mm-Tap1-C2 | ACD Biotechnne | 575041-C2 | NA | FISH |
| Pan-MHC Class 1** | ACD Biotechnne | NPR-0023100 | NA | FISH |
| TSA Cy3 | Perkin Elmer | NEL744001KT | NA | FISH, 1:1500 |
| TSA Cy5 | Perkin Elmer | NEL745001KT | NA | FISH, 1:1500<br>IHC, 1:7500 |
| Recombinant Mouse IFN-gamma Protein | R&D Systems<br>(Biotechnne) | 485-MI-100/CF | NA | 100 ng/μL |

FANS: fluorescence-activated nuclear sorting, IHC: immunohistochemical staining of brain sections, FISH: Fluorescence *in situ* hybridization.
\*\*Due to extensive homology between MHC Class I gene family members, probes selectively targeting *H2-K1* and/or *H2-D1* were not feasible. The new probe employed was designed to maximally cross-react with *H2-K1* mRNA (NCBI Reference Sequence, NM\_001001892.2), although substantial cross-reactivity is predicted for *H2-K2*, *H2-D1*, *H2-D4*, *H2-Q7*, *H2-Q9*, *H2-Q10*, and *Gm38417*.

### Immunohistochemistry (IHC)

Mice were anesthetized with euthasol and transcardially perfused with ∼15 mL of 0.9% saline followed by 40-50 mL of ice-cold 4% paraformaldehyde (PFA) in 0.1 M phosphate buffer (PB), pH 7.4. Brains were post-fixed in 4% PFA in 0.1M PB for 6-12 hours at 4°C, washed three times in phosphate buffered saline (PBS), and sectioned at 50 μm on a Leica VT1000S vibratome. Sections were placed in cryoprotectant solution (30% ethylene glycol, 30% glycerol, 0.1M PB, pH 7.4) and stored at −20°C until further use.

Sections were removed from cryoprotectant solution and washed three times in tris-buffered saline (TBS) at room temperature. Except for pSTAT1 staining, sections were then permeabilized in TBS + 0.2% Triton-X 100 for one hour at room temperature, followed by blocking in TBS + 10% normal goat serum (NGS) and 0.1% Triton-X 100 for 1.5 hours at room temperature. Sections for pSTAT1 staining were slide mounted, submerged in 10 mM sodium citrate (pH 6.0) at 50°C for 15 minutes, and washed several times in TBS. All sections were then transferred to or submerged in a pre-chilled solution containing primary antibodies in TBS + 2% NGS + 0.1% Triton-X 100 and incubated overnight at 4°C. Sections were washed in TBS + 0.05% Tween 20 (TBS+T) five times over an hour at room temperature. Sections were incubated in a solution containing secondary antibodies in TBS + 2% NGS + 0.1% Triton-X 100 at room temperature for 1.5 hours, followed by four washes in TBS+T over 45 minutes at room temperature. Sections were slide mounted and coverslipped with Fluoromount G (Southern Biotech). See *Antibodies and Reagents* for a complete list of antibodies and concentrations used in this study.

### Fluorescence in situ hybridization (FISH)

FISH was performed using the highly sensitive RNAScope® Multiplex Fluorescent v2 assay (ACD Bio). See *Antibodies and Reagents* for complete list of probes and reagents used in this study. Although most single FISH puncta using this assay are likely single mRNA molecules (Wang et al., 2012), this cannot be definitively determined due to the enzymatic signal amplification and non-diffraction-limited size of the mRNA puncta.

Mouse brain sections were prepared as above, removed from cryoprotectant solution, and washed three times in tris-buffered saline (TBS) at room temperature. Sections were incubated with hydrogen peroxide (ACD) for 15 minutes at room temperature, washed several times in TBS, and then mounted to Superfrost slides (Fisher).

Sections were allowed to dry for 10 minutes and a hydrophobic barrier (PAP pen, Vector Labs) was created around the tissue. Tissue was incubated in 50% EtOH, then 70% EtOH, then 100% EtOH for 5 minutes each. Sections were rehydrated in TBS for several minutes, digested with Protease IV (ACD) for 25 minutes at room temperature, and rinsed twice with TBS before proceeding to the RNA Scope Multiplex Fluorescent v2 assay (ACD).

The RNA Scope Multiplex Fluorescent v2 assay was conducted according to the manufacturer’s instructions, with all incubations taking place in a humidified chamber at 40°C. Two 5-minute washes in excess RNA Scope Wash Buffer (ACD) took place between each incubation in sequential order: probes (2-hours), AMP1 (30 minutes), AMP2 (30 minutes), AMP3 (15 minutes), HRP-C1/2/3 (15 minutes), TSA Cy3 (30 minutes), HRP blocker (30 minutes), HRP-C1/2/3 (15 minutes), and TSA Cy5 (30 minutes). Samples were washed twice more in RNA Scope Wash Buffer, then twice more in TBS. Samples were then blocked and immunostained for Tyrosine Hydroxylase as described above. After immunostaining, samples were mounted in Fluoromount G and stored at 4°C for up to 1 week before imaging.

### Image acquisition and analysis

All imaging of 50 μm sections from perfusion-fixed brain was conducted either on a Nikon Ti2 Eclipse epifluorescence microscope or on a Leica SP8 scanning confocal microscope using a 20x/0.75 NA or 60x/1.4 NA objective.

For pStat1-Y701 analysis, 5 μm Z stacks from the SNc were acquired using a 20x/0.75 NA objective and collapsed via maximum projection. Three types of DAPI^+^ nuclei were segmented as ROIs: TH+/NeuN+ (mDA neurons), TH-/NeuN+ (non mDA neurons), and glia (TH-/NeuN-). The mean pStat1 intensity for all pixels within each ROI was background subtracted using the median of the entire image.

For FISH analysis, 5 μm Z stacks from the SNc were acquired using a 20x/0.75 NA objective and collapsed via maximum projection. For quantification of mRNA density in neuronal somata, a binary threshold was set for the TH immunofluorescence signal based on two standard deviations above the image background to generate a binary mask of pixels for TH^+^ neurons. Segmentation of mDA neuronal somata was conducted in ImageJ 1.53c (RRID:SCR_003070, https://imagej.net/) to generate ROIs for each neuronal soma, after which the total signal in both RNA channels was measured. The mean RNA intensity for all pixels within each soma ROI was background subtracted using the median of the entire image.

### Fluorescence-Activated Nuclei Sorting (FANS)

Nuclei were prepared from the ventral midbrain of mice at 48 hours after injection of saline or IFN-γ. Mice were sacrificed by cervical dislocation and brains were rapidly extracted and submerged in ice-cold phosphate-buffered saline (PBS). Brains were placed on an ice-cold brain matrix (Zivic Instruments) and separated into 0.5-1.0 mm sections using ice cold razor blades. Ventral midbrain tissue was dissected from slices between approximately -2.5 mm to -3.75 mm AP to Bregma. First, the cortex, hippocampi, and any hypothalamus or white matter ventral to the midbrain were removed. A single horizontal cut was made just dorsal to the rostral linear nucleus and all dorsal tissue was discarded. The remaining tissue containing the SN/VTA was flash frozen on liquid nitrogen and stored at-80°C.

Nuclei preparation and sorting were conducted as described by Krishnaswami et al. (2016) with only minor modifications. Frozen tissue was thawed in nuclear isolation medium 1 (NIM1: 250 mM sucrose, 25 mM KCl, 5 mM MgCl_2_, 10 mM Tris, pH 8.0) supplemented with 1 mM DTT, 0.2 U/μL SUPERaseIN, 1x EDTA-free Protease Inhibitor Cocktail (Roche), and 0.1% Triton X-100. Tissue was homogenized on ice in a glass-glass dounce homogenizer with 30 gentle strokes of loose and tight clearance pestles. All subsequent purification steps were performed on ice or at 4°C unless otherwise specified. Lysates were spun at 500xg for 5 minutes, resuspended in NIM1 supplemented as above and also with 1% BSA, and passed through a 40μm cell strainer cap. Nuclei were centrifuged again at 500xg for 5 minutes and resuspended in staining buffer (PBS supplemented with 1% BSA, 5 mM MgCl_2_, 0.2 U/μL SUPERaseIN, 1 μg/mL propidium iodide, and 2 μg/mL mouse anti-NeuN AlexaFluor488). After 30 minutes incubation on ice, nuclei were centrifuged at 500xg for 5 minutes and resuspended in 0.5 mL of FACS buffer (PBS + 0.04% BSA, 0.2 U/μL SUPERaseIN, and 1 μg/mL propidium iodide) for sorting.

Nuclei sorting was conducted on a BD Influx (BD Biosciences) using a 100 μm nozzle at 11.1 psi. Single nuclei were gated first using FSC-H vs. FSC-A and then on propidium iodide fluorescence. NeuN/AlexaFluor488 fluorescence was used to establish NeuN^-^ and NeuN^+^ gates on the single nuclei population. Nuclei were sorted directly into 96 well plates (for bulk nuclei RNA-seq with 96-well plate, pooled library construction) or into 1.5 mL Eppendorf tubes that had been previously coated with 0.1% BSA in PBS overnight at 37°C (for snRNA-seq).

#### Low input RNA Sequencing with 96-well plate, pooled library construction

The protocol for plate based, 3’ end unique molecular indicator (UMI)-based RNA sequencing of single cells has been described previously (Snyder et al., 2019) as well as modifications to accommodate ultra-low input samples (Hobson et al., 2022b). See **Supplementary File 1** for sequences of all custom primers and oligonucleotides used in this protocol. For neuronal and glial nuclei analysis in **Figure 1-2**, 100 NeuN^-^ or NeuN^+^ nuclei were sorted directly into the wells of a 96 well plate containing 3 μL of nuclease-free water containing 1 U/μL SUPERaseIN (sort volume ∼3 μL for total volume of ∼6 μL). After adding 1.5 μL of 10 μM barcoded RT primer (Integrated DNA Technologies), primer annealing was performed at 72°C for 3 minutes. Reverse transcription was performed by adding 7.5 μL RT mix to each well (2.81 μL of 40% polyethylene glycol 8000, 0.15 μL of 100 mM dNTPs, 3 μL of 5X Maxima H RT Buffer, 0.2 μL of 200 U/μL Maxima H Reverse Transcriptase (ThermoFisher), 0.2 μL of 20 U/μL SUPERaseIN, and 0.15 μL of 100 μM Template Switching Oligo (Integrated DNA Technologies), and 1 μL of nuclease free water). Reverse transcription was performed at 42°C for 90 minutes, followed by 10 cycles of 50°C for 2 minutes, 42°C for 2 minutes, 75°C for 10 minutes, followed by a 4°C hold. Excess primers were removed by adding 2 μL of Exonuclease I mix (1.875U ExoI in water) to each well and incubating at 37°C for 30 minutes, 85°C for 15 minutes, 75°C for 30 seconds, 4°C hold.

All wells were pooled into a single 15-ml falcon tubes and cDNA was purified and concentrated using Dynabeads^™^ MyOne^™^ Silane beads (ThermoFisher) according to the manufacturer’s instructions. The cDNA was split into duplicate reactions containing 25 μl cDNA, 25 μl 2x HIFI HotStart Ready Mix (Kapa Biosystems), and 0.2 M SMART PCR Primer. PCR was run as follows: 37°C for 30 minutes, 85°C for 15 minutes, 75°C for 30 seconds, 4°C hold. Duplicate reactions were combined and purified using 0.7 volumes AMPure XP beads (Beckman Coulter). The amplified cDNA was visualized on an Agilent 2100 Bioanalyzer and quantified using a Qubit II fluorometer (ThermoFisher).

Sequencing libraries were constructed using Nextera XT (Illumina) with modifications. A custom i5 primer was used (NexteraPCR) with 0.6ng input cDNA and 10 cycles of amplification was performed. Unique i7 indexes were used for each plate. After amplification, the library was purified with two rounds of AMPure XP beads, visualized on the Agilent 2100 Bioanalyzer and quantified using the Qubit II fluorometer. Libraries were sequenced on an Illumina NextSeq 500 using the 75 cycle High Output kit (read lengths 26(R1) x 8(i) x 58(R2)). Custom sequencing primers were used for Read 1 (SMRT_R1seq and ILMN_R1seq, see *Antibodies and Reagents*). With each plate we targeted ∼400 M reads. Library pools were loaded at 1.8 pM with 20% PhiX (Illumina).

Reads were aligned to the mouse reference genome GRCm38 and transcriptome annotation (Gencode vM10) using the STAR aligner with parameters *–sjdbOverhang 65 –twopassMode Basic* after trimming poly(A)-tails from the 3’-ends. The aligned reads were demultiplexed using the well-identifying barcodes, correcting all single-nucleotide errors. All reads with the same well-identifying barcode, UMI, and gene mapping were collapsed to represent an individual transcript. To correct for sequencing errors in UMIs, we further collapsed UMIs that were within Hamming distance one of another UMI with the same well-identifying barcode and gene. For each 96-well plate, after generating a final list of individual transcripts with unique combinations of well-identifying barcodes, UMIs, and gene mapping, we produced a molecular count matrix for downstream analysis.

### Bulk Nuclei RNA-Seq Differential Expression Analysis

Analysis of bulk nuclei RNA-Seq UMI count matrices as shown in **Figure 1-2** (from *Low input RNA Sequencing with 96-well plate, pooled library construction*) was conducted using a generalized linear model (GLM) in *DESeq2* (Love et al., 2014). For analysis of IFN-γ vs. saline analyses, a single contrast was made for each subset of NeuN-or NeuN+ nuclei at each timepoint (*DESeq2* formula: ∼Treatment). For interaction analysis at each timepoint (*DESeq2* formula: ∼NeuN + Treatment + NeuN:Treatment), the contrast specifies the difference in log2 Fold Change for the effect of Treatment (IFN-γ vs. saline) between the levels of NeuN (i.e., for NeuPos.IFN, the difference in IFN-γ vs. saline between NeuN- and NeuN+ samples). Global analysis of NeuN-vs. NeuN+ nuclei was conducted across all samples while controlling for the effect of IFN-γ (*DESeq2* formula: ∼Treatment + NeuN). Complete *DESeq2* output for all analyses related to **Figure 1** and **Figure 2** is found in **Supplementary Table 1.**

### Single nucleus RNA-sequencing (snRNA-seq)

NeuN nuclear sorting was conducted as described above. Approximately 70% NeuN^+^ and 30% NeuN^-^ single nuclei were sorted per sample. Sorted nuclei were placed on ice and immediately submitted to the Single Cell Analysis Core (Columbia Genome Center) for 10X Genomics Chromium Single Cell 3’ v3 library preparation. The resulting libraries were sequenced on an Illumina NovaSeq 6000 with a targeted depth of >100,000 2×100 bp paired-end reads per nucleus.

### snRNA-seq Data Processing, Clustering, and Doublet Filtering

snRNA-seq data were processed as described previously (Yuan et al., 2018) (code available at: https://github.com/simslab/DropSeqPipeline8). Reads were aligned to the mouse reference genome GRCm38 and transcriptome annotation (Gencode vM10) using the STAR aligner with parameters *–sjdbOverhang 65 – twopassMode Basic* after trimming poly(A)-tails from the 3’-ends. All reads that uniquely aligned to a given gene body (including introns) were kept for downstream filtering analysis, because a large fraction of nuclear transcripts are unspliced. The aligned reads were demultiplexed using the cell-identifying barcodes, correcting all single-nucleotide errors. All reads with the same cell-identifying barcode, UMI, and gene mapping were collapsed to represent an individual transcript. To correct for sequencing errors in UMIs, we further collapsed UMIs that were within Hamming distance one of another UMI with the same cell-identifying barcode and gene. For each sample, after generating a final list of individual transcripts with unique combinations of cell-identifying barcodes, UMIs, and gene mapping, we produced a molecular count matrix for downstream analysis.

We first performed unsupervised clustering on the count matrices using the PhenoGraph v1.5.7 (Levine et al., 2015) (RRID:SCR_016919, https://github.com/dpeerlab/phenograph) implementation of Louvain community detection after selection of highly variable genes and construction of a k-nearest neighbors graph as described previously (Levitin et al., 2019). We identified 38 clusters (**Supp. Fig. 4a**), two of which showed statistically significant co-enrichment of dopamine neuron markers such as *Th* and *Slc6a3* based on the binomial test for expression specificity (Shekhar et al., 2016). We identified six major cell types with sufficient coverage for downstream analysis as shown in the UMAP (Becht et al., 2018) embedding in **Figure 3a**. The assignment of PhenoGraph clusters into major cell type groupings was based on key marker genes significantly enriched in each cluster, as shown in **Supp. Fig. 4b:** Oligodendrocytes (*Mal, Mag, Mog*), Astrocytes (*Gja1, Fgfr3, Atp1a2*), Microglia (*C1qb, Siglech, Csf1r*), VGLUT2 neurons (*Slc17a6, Cacna2d1, Ntng1*), GABA neurons (*Slc32a1, Gad1, Gad2*), and DA neurons (*Th, Slc6a3, En1, Slc10a4*). Rare populations with insufficient cell numbers (e.g., mural cells, T cells) or ambiguous neuronal clusters possibly containing multiplets (e.g., Gad1^+^/Gad2^+^/Slc17a7^+^/Th^+^) were excluded from downstream analysis.

While examining the neuronal profiles, we identified small subpopulations of nuclei with co-enrichment of neuronal and oligodendrocyte markers such as *Mbp, Plp1, Mog*, and *Mag*. To filter out oligodendrocyte doublets, we fit Gaussian mixture models to the sum of expression for oligodendrocyte marker genes within each major neuronal class (see **Supp. Fig. 4c**). The top 99 oligodendrocyte marker genes were used, as determined by the binomial test (log2 specificity > 3, FDR < 10e-50). For all non-oligodendrocyte clusters, we then removed nuclei with expression >8 standard deviations above the mean of the first Gaussian component corresponding to single neurons (**Supp. Fig. 4d**). This process was repeated twice more using the top 99 astrocyte and top 99 microglial marker genes to filter out potential doublets.

### snRNA-Seq Differential Expression Analysis

For differential expression analysis shown in **Fig. 3**, we identified genes upregulated or downregulated by IFN-γ within each cell type after filtering by conducting differential expression analysis as described in (Zhao et al., 2021). Briefly, to perform differential expression analysis between two groups of cells, we randomly sub-sampled the data so that both groups are represented by the same number of cells. Next, we randomly sub-sampled the detected transcripts so that both groups have the same average number of transcripts per cell. Finally, we normalized the two sub-sampled count matrices using *scran* (Lun et al., 2016) and analyzed differential expression for each gene using the SciPy implementation of the Mann-Whitney U-test. We corrected the resulting p-values for false discovery using the Benjamini-Hochberg procedure as implemented in the *statsmodels 0.13.4* (RRID:SCR_016074, http://www.statsmodels.org/) package in Python 3.7 (Python Programming Language, RRID:SCR_008394, http://www.python.org/). We compared each IFN-γ sample to the saline control and retained only genes with >8-fold differential expression |log_2_FC > 3| and FDR < 0.01 in both IFN-γ samples. Complete differential expression output for all analyses related to **Figure 3** is found in **Supplementary Table 2.**

### Gene Ontology (GO) Analysis

A single list of unique genes was used (i.e., differentially expressed genes from *DESeq2* analysis) to conduct GO Analysis using web-based Enrichr v3.29.21 (Xie et al., 2021) (RRID:SCR_001575, http://amp.pharm.mssm.edu/Enrichr/) with 2018 GO Terms for Cellular Component, Biological Process, and Molecular Function (Ashburner et al., 2000; Gene Ontology Consortium, 2021).

### Visualization and Statistical Analysis

Graphics were created in Adobe Illustrator 24.3 (Adobe Inc., http://www.adobe.com/products/illustrator.html, RRID:SCR_010279). Unless otherwise noted, all statistical analysis and data visualization was conducted in Python using *SciPy 1.9.2* (RRID:SCR_008058, http://www.scipy.org/), *Matplotlib 3.6.0* (RRID:SCR_008624, http://matplotlib.sourceforge.net/), and *Seaborn 0.12.1* (RRID:SCR_018132, https://seaborn.pydata.org/) packages. Statistical comparisons were conducted using the Mann-Whitney U test with number of replicates and other statistical testing information indicated in the figure captions. For visualization of RNA-seq abundance data, mRNA UMI counts were scaled up by 10^6^ and normalized to total counts (CPM), and log2 transformed after adding 1. Normalized mRNA abundance is referred to as ‘log2(CPM+1)’ as specified in figure captions. For clustered heatmaps, Z-scores of log2(CPM+1) were first calculated using the *zscore* function within the *SciPy Stats* module, after which the row and column clustering was calculated using the *linkage* function (metric = ‘Euclidean’, method= ‘average’) within *fastcluster 1.2.3* (Müllner, 2013) (https://github.com/dmuellner/fastcluster) and passed to *Seaborn clustermap*.

## Funding

This work was conducted in collaboration with the JP Sulzberger Columbia Genome Center at Columbia University Irving Medical Center. This research was funded in part by Aligning Science Across Parkinson’s [ASAP-000375] (DS and PAS) through the Michael J. Fox Foundation for Parkinson’s Research (MJFF). For the purpose of open access, the author has applied a CC BY public copyright license to all Author Accepted Manuscripts arising from this submission. This work was supported by the JPB Foundation (DS). This work was supported by NIH grants F30 DA047775-04 (BDH), R01 NS0954 (DS), R01 DA07418 (DS), and R01 MH122470 (DS). We thank Vanessa Morales for assistance with animal colony management.

## Author Contributions

BDH, DS, and PAS conceived the overall project. ATS and BDH executed stereotaxic injections. BDH executed tissue dissection, fluorescence activated nuclear sorting, low input bulk RNA-seq, and snRNA-seq experiments. BDH conducted bulk nuclear RNA-seq analysis. BDH, BC, and PAS conducted snRNA-seq analysis. BDH executed IHC experiments with assistance from EVM. BDH and MBD conducted FISH experiments. BDH analyzed IHC and FISH imaging data. PAS and DS supervised the research. BDH wrote the manuscript with input from PAS and DS. All authors edited, read, and approved the final manuscript.

## Competing interests

The authors declare that no competing interests exist.

## Data and materials availability

There are restrictions to the availability of custom-order RNA Scope probes generated for this study due to limited production size. The exact new probe design IDs are listed in *Antibodies and Reagents* and are available for further production from ACD Biotechne.

All bulk nuclear RNA-seq and snRNA-seq data from this study are publicly available on the Gene Expression Omnibus as GSE218132. Count matrices from this study are in **Supplementary File 3** and **Supplementary File 5**.

Python and Shell code used for processing of RNA-seq data is accessible at: https://github.com/simslab/DropSeqPipeline8, and Python code used for clustering and visualization of scRNA-seq data can be found at www.github.com/simslab/cluster_diffex2018.

## Notes

https://www.ncbi.nlm.nih.gov/geo/query/acc.cgi?acc=GSE218132

### Competing Interest Statement

The authors have declared no competing interest.

### Summary of Updates

some pdf conversion line errors corrected order of authors corrected

## References

Amor, S., Peferoen, L.A.N., Vogel, D.Y.S., Breur, M., van der Valk, P., Baker, D., van Noort, J.M., 2014. Inflammation in neurodegenerative diseases--an update. Immunology 142, 151–166. https://doi.org/10.1111/imm.12233

Ashburner, M., Ball, C.A., Blake, J.A., Botstein, D., Butler, H., Cherry, J.M., Davis, A.P., Dolinski, K., Dwight, S.S., Eppig, J.T., Harris, M.A., Hill, D.P., Issel-Tarver, L., Kasarskis, A., Lewis, S., Matese, J.C., Richardson, J.E., Ringwald, M., Rubin, G.M., Sherlock, G., 2000. Gene Ontology: tool for the unification of biology. Nat Genet 25, 25–29. https://doi.org/10.1038/75556

Babcock, A.A., Kuziel, W.A., Rivest, S., Owens, T., 2003. Chemokine expression by glial cells directs leukocytes to sites of axonal injury in the CNS. J Neurosci 23, 7922–7930.

Bäckman, C.M., Malik, N., Zhang, Y., Shan, L., Grinberg, A., Hoffer, B.J., Westphal, H., Tomac, A.C., 2006. Characterization of a mouse strain expressing Cre recombinase from the 3’ untranslated region of the dopamine transporter locus. Genesis 44, 383–390. https://doi.org/10.1002/dvg.20228

Barcia, C., Ros, C.M., Annese, V., Gómez, A., Ros-Bernal, F., Aguado-Yera, D., Martínez-Pagán, M.E., de Pablos, V., Fernandez-Villalba, E., Herrero, M.T., 2011. IFN-γ signaling, with the synergistic contribution of TNF-α, mediates cell specific microglial and astroglial activation in experimental models of Parkinson’s disease. Cell Death Dis 2, e142. https://doi.org/10.1038/cddis.2011.17

Becht, E., McInnes, L., Healy, J., Dutertre, C.-A., Kwok, I.W.H., Ng, L.G., Ginhoux, F., Newell, E.W., 2018. Dimensionality reduction for visualizing single-cell data using UMAP. Nat Biotechnol. https://doi.org/10.1038/nbt.4314

Bernard-Valnet, R., Yshii, L., Quériault, C., Nguyen, X.-H., Arthaud, S., Rodrigues, M., Canivet, A., Morel, A.-L., Matthys, A., Bauer, J., Pignolet, B., Dauvilliers, Y., Peyron, C., Liblau, R.S., 2016. CD8 T cell-mediated killing of orexinergic neurons induces a narcolepsy-like phenotype in mice. Proc. Natl. Acad. Sci. U.S.A. 113, 10956–10961. https://doi.org/10.1073/pnas.1603325113

Brochard, V., Combadière, B., Prigent, A., Laouar, Y., Perrin, A., Beray-Berthat, V., Bonduelle, O., Alvarez-Fischer, D., Callebert, J., Launay, J.-M., Duyckaerts, C., Flavell, R.A., Hirsch, E.C., Hunot, S., 2009. Infiltration of CD4+ lymphocytes into the brain contributes to neurodegeneration in a mouse model of Parkinson disease. J Clin Invest 119, 182–192. https://doi.org/10.1172/JCI36470

Cebrián, C., Zucca, F.A., Mauri, P., Steinbeck, J.A., Studer, L., Scherzer, C.R., Kanter, E., Budhu, S., Mandelbaum, J., Vonsattel, J.P., Zecca, L., Loike, J.D., Sulzer, D., 2014. MHC-I expression renders catecholaminergic neurons susceptible to T-cell-mediated degeneration. Nat Commun 5, 3633. https://doi.org/10.1038/ncomms4633

Chakrabarty, P., Ceballos-Diaz, C., Lin, W.-L., Beccard, A., Jansen-West, K., McFarland, N.R., Janus, C., Dickson, D., Das, P., Golde, T.E., 2011. Interferon-γ induces progressive nigrostriatal degeneration and basal ganglia calcification. Nat Neurosci 14, 694–696. https://doi.org/10.1038/nn.2829

Chung, C.Y., Seo, H., Sonntag, K.C., Brooks, A., Lin, L., Isacson, O., 2005. Cell type-specific gene expression of midbrain dopaminergic neurons reveals molecules involved in their vulnerability and protection. Hum Mol Genet 14, 1709– 1725. https://doi.org/10.1093/hmg/ddi178

Clarkson, B.D.S., Patel, M.S., LaFrance-Corey, R.G., Howe, C.L., 2018. Retrograde interferon-gamma signaling induces major histocompatibility class I expression in human-induced pluripotent stem cell-derived neurons. Ann Clin Transl Neurol 5, 172–185. https://doi.org/10.1002/acn3.516

Corriveau, R.A., Huh, G.S., Shatz, C.J., 1998. Regulation of Class I MHC Gene Expression in the Developing and Mature CNS by Neural Activity. Neuron 21, 505–520. https://doi.org/10.1016/S0896-6273(00)80562-0

Dhanwani, R., Lima-Junior, J.R., Sethi, A., Pham, J., Williams, G., Frazier, A., Xu, Y., Amara, A.W., Standaert, D.G., Goldman, J.G., Litvan, I., Alcalay, R.N., Peters, B., Sulzer, D., Arlehamn, C.S.L., Sette, A., 2022. Transcriptional analysis of peripheral memory T cells reveals Parkinson’s disease-specific gene signatures. npj Parkinsons Dis. 8, 1– 10. https://doi.org/10.1038/s41531-022-00282-2

Di Liberto, G., Pantelyushin, S., Kreutzfeldt, M., Page, N., Musardo, S., Coras, R., Steinbach, K., Vincenti, I., Klimek, B., Lingner, T., Salinas, G., Lin-Marq, N., Staszewski, O., Costa Jordão, M.J., Wagner, I., Egervari, K., Mack, M., Bellone, C., Blümcke, I., Prinz, M., Pinschewer, D.D., Merkler, D., 2018. Neurons under T Cell Attack Coordinate Phagocyte-Mediated Synaptic Stripping. Cell 175, 458-471.e19. https://doi.org/10.1016/j.cell.2018.07.049

Dulken, B.W., Buckley, M.T., Navarro Negredo, P., Saligrama, N., Cayrol, R., Leeman, D.S., George, B.M., Boutet, S.C., Hebestreit, K., Pluvinage, J.V., Wyss-Coray, T., Weissman, I.L., Vogel, H., Davis, M.M., Brunet, A., 2019. Single-cell analysis reveals T cell infiltration in old neurogenic niches. Nature 571, 205–210. https://doi.org/10.1038/s41586-019-1362-5

Falcão, A.M., van Bruggen, D., Marques, S., Meijer, M., Jäkel, S., Agirre, E., Samudyata, null, Floriddia, E.M., Vanichkina, D.P., Ffrench-Constant, C., Williams, A., Guerreiro-Cacais, A.O., Castelo-Branco, G., 2018. Disease-specific oligodendrocyte lineage cells arise in multiple sclerosis. Nat Med 24, 1837–1844. https://doi.org/10.1038/s41591-018-0236-y

Ferrari, D.P., Bortolanza, M., Del Bel, E.A., 2021. Interferon-γ Involvement in the Neuroinflammation Associated with Parkinson’s Disease and L-DOPA-Induced Dyskinesia. Neurotox Res 39, 705–719. https://doi.org/10.1007/s12640-021-00345-x

Filiano, A.J., Xu, Y., Tustison, N.J., Marsh, R.L., Baker, W., Smirnov, I., Overall, C.C., Gadani, S.P., Turner, S.D., Weng, Z., Peerzade, S.N., Chen, H., Lee, K.S., Scott, M.M., Beenhakker, M.P., Litvak, V., Kipnis, J., 2016. Unexpected role of interferon-γ in regulating neuronal connectivity and social behavior. Nature 535, 425–429. https://doi.org/10.1038/nature18626

Flood, L., Korol, S.V., Ekselius, L., Birnir, B., Jin, Z., 2019. Interferon-γ potentiates GABAA receptor-mediated inhibitory currents in rat hippocampal CA1 pyramidal neurons. J Neuroimmunol 337, 577050. https://doi.org/10.1016/j.jneuroim.2019.577050

Galiano-Landeira, J., Torra, A., Vila, M., Bové, J., 2020. CD8 T cell nigral infiltration precedes synucleinopathy in early stages of Parkinson’s disease. Brain 143, 3717–3733. https://doi.org/10.1093/brain/awaa269

Garretti, F., Monahan, C., Sette, A., Agalliu, D., Sulzer, D., 2022. T cells, α-synuclein and Parkinson disease. Handb Clin Neurol 184, 439–455. https://doi.org/10.1016/B978-0-12-819410-2.00023-0

Gate, D., Saligrama, N., Leventhal, O., Yang, A.C., Unger, M.S., Middeldorp, J., Chen, K., Lehallier, B., Channappa, D., De Los Santos, M.B., McBride, A., Pluvinage, J., Elahi, F., Tam, G.K.-Y., Kim, Y., Greicius, M., Wagner, A.D., Aigner, L., Galasko, D.R., Davis, M.M., Wyss-Coray, T., 2020. Clonally expanded CD8 T cells patrol the cerebrospinal fluid in Alzheimer’s disease. Nature 577, 399–404. https://doi.org/10.1038/s41586-019-1895-7

Gate, D., Tapp, E., Leventhal, O., Shahid, M., Nonninger, T.J., Yang, A.C., Strempfl, K., Unger, M.S., Fehlmann, T., Oh, H., Channappa, D., Henderson, V.W., Keller, A., Aigner, L., Galasko, D.R., Davis, M.M., Poston, K.L., Wyss-Coray, T., 2021. CD4+ T cells contribute to neurodegeneration in Lewy body dementia. Science 374, 868–874. https://doi.org/10.1126/science.abf7266

Gene Ontology Consortium, 2021. The Gene Ontology resource: enriching a GOld mine. Nucleic Acids Res 49, D325–D334. https://doi.org/10.1093/nar/gkaa1113

Gottfried-Blackmore, A., Kaunzner, U.W., Idoyaga, J., Felger, J.C., McEwen, B.S., Bulloch, K., 2009. Acute in vivo exposure to interferon-gamma enables resident brain dendritic cells to become effective antigen presenting cells. Proc Natl Acad Sci U S A 106, 20918–20923. https://doi.org/10.1073/pnas.0911509106

Harrison, J.K., Jiang, Y., Chen, S., Xia, Y., Maciejewski, D., McNamara, R.K., Streit, W.J., Salafranca, M.N., Adhikari, S., Thompson, D.A., Botti, P., Bacon, K.B., Feng, L., 1998. Role for neuronally derived fractalkine in mediating interactions between neurons and CX3CR1-expressing microglia. Proc Natl Acad Sci U S A 95, 10896–10901. https://doi.org/10.1073/pnas.95.18.10896

Hirsch, E., Graybiel, A.M., Agid, Y.A., 1988. Melanized dopaminergic neurons are differentially susceptible to degeneration in Parkinson’s disease. Nature 334, 345–348. https://doi.org/10.1038/334345a0

Hobson, B.D., Choi, S.J., Mosharov, E.V., Soni, R.K., Sulzer, D., Sims, P., 2022a. Subcellular proteomics of dopamine neurons in the mouse brain. Elife 11, e70921. https://doi.org/10.7554/eLife.70921

Hobson, B.D., Kong, L., Angelo, M.F., Lieberman, O.J., Mosharov, E.V., Herzog, E., Sulzer, D., Sims, P.A., 2022b. Subcellular and regional localization of mRNA translation in midbrain dopamine neurons. Cell Rep 38, 110208. https://doi.org/10.1016/j.celrep.2021.110208

Höftberger, R., Aboul-Enein, F., Brueck, W., Lucchinetti, C., Rodriguez, M., Schmidbauer, M., Jellinger, K., Lassmann, H., 2004. Expression of major histocompatibility complex class I molecules on the different cell types in multiple sclerosis lesions. Brain Pathol 14, 43–50. https://doi.org/10.1111/j.1750-3639.2004.tb00496.x

Howcroft, T.K., Singer, D.S., 2003. Expression of nonclassical MHC class Ib genes: comparison of regulatory elements. Immunol Res 27, 1–30. https://doi.org/10.1385/IR:27:1:1

Howe, C.L., LaFrance-Corey, R.G., Goddery, E.N., Johnson, R.K., Mirchia, K., 2017. Neuronal CCL2 expression drives inflammatory monocyte infiltration into the brain during acute virus infection. J Neuroinflammation 14, 238. https://doi.org/10.1186/s12974-017-1015-2

Isaacs, A., Lindenmann, J., Andrewes, C.H., 1957. Virus interference. I. The interferon. Proceedings of the Royal Society of London. Series B - Biological Sciences 147, 258–267. https://doi.org/10.1098/rspb.1957.0048

Ivashkiv, L.B., 2018. IFNγ: signalling, epigenetics and roles in immunity, metabolism, disease and cancer immunotherapy. Nat Rev Immunol 18, 545–558. https://doi.org/10.1038/s41577-018-0029-z

Iwata, H., Goettsch, C., Sharma, A., Ricchiuto, P., Goh, W.W.B., Halu, A., Yamada, I., Yoshida, H., Hara, T., Wei, M., Inoue, N., Fukuda, D., Mojcher, A., Mattson, P.C., Barabási, A.-L., Boothby, M., Aikawa, E., Singh, S.A., Aikawa, M., 2016. PARP9 and PARP14 cross-regulate macrophage activation via STAT1 ADP-ribosylation. Nat Commun 7, 12849. https://doi.org/10.1038/ncomms12849

Jäkel, S., Agirre, E., Mendanha Falcão, A., van Bruggen, D., Lee, K.W., Knuesel, I., Malhotra, D., ffrench-Constant, C., Williams, A., Castelo-Branco, G., 2019. Altered human oligodendrocyte heterogeneity in multiple sclerosis. Nature 566, 543–547. https://doi.org/10.1038/s41586-019-0903-2

Jongsma, M.L.M., Guarda, G., Spaapen, R.M., 2019. The regulatory network behind MHC class I expression. Mol Immunol 113, 16–21. https://doi.org/10.1016/j.molimm.2017.12.005

Kaya, T., Mattugini, N., Liu, L., Ji, H., Cantuti-Castelvetri, L., Wu, J., Schifferer, M., Groh, J., Martini, R., Besson-Girard, S., Kaji, S., Liesz, A., Gokce, O., Simons, M., 2022. CD8+ T cells induce interferon-responsive oligodendrocytes and microglia in white matter aging. Nat Neurosci 1–12. https://doi.org/10.1038/s41593-022-01183-6

Kreutzfeldt, M., Bergthaler, A., Fernandez, M., Brück, W., Steinbach, K., Vorm, M., Coras, R., Blümcke, I., Bonilla, W.V., Fleige, A., Forman, R., Müller, W., Becher, B., Misgeld, T., Kerschensteiner, M., Pinschewer, D.D., Merkler, D., 2013. Neuroprotective intervention by interferon-γ blockade prevents CD8+ T cell–mediated dendrite and synapse loss. J Exp Med 210, 2087–2103. https://doi.org/10.1084/jem.20122143

Krishnaswami, S.R., Grindberg, R.V., Novotny, M., Venepally, P., Lacar, B., Bhutani, K., Linker, S.B., Pham, S., Erwin, J.A., Miller, J.A., Hodge, R., McCarthy, J.K., Kelder, M., McCorrison, J., Aevermann, B.D., Fuertes, F.D., Scheuermann, R.H., Lee, J., Lein, E.S., Schork, N., McConnell, M.J., Gage, F.H., Lasken, R.S., 2016. Using single nuclei for RNA-seq to capture the transcriptome of postmortem neurons. Nat Protoc 11, 499–524. https://doi.org/10.1038/nprot.2016.015

Kustrimovic, N., Comi, C., Magistrelli, L., Rasini, E., Legnaro, M., Bombelli, R., Aleksic, I., Blandini, F., Minafra, B., Riboldazzi, G., Sturchio, A., Mauri, M., Bono, G., Marino, F., Cosentino, M., 2018. Parkinson’s disease patients have a complex phenotypic and functional Th1 bias: cross-sectional studies of CD4+ Th1/Th2/T17 and Treg in drug-naïve and drug-treated patients. Journal of Neuroinflammation 15, 205. https://doi.org/10.1186/s12974-018-1248-8

Lee, S.H., Carrero, J.A., Uppaluri, R., White, J.M., Archambault, J.M., Lai, K.S., Chan, S.R., Sheehan, K.C.F., Unanue, E.R., Schreiber, R.D., 2013. Identifying the initiating events of anti-Listeria responses using mice with conditional loss of IFN-γ receptor subunit 1 (IFNGR1). J. Immunol. 191, 4223–4234. https://doi.org/10.4049/jimmunol.1300910

Levine, J.H., Simonds, E.F., Bendall, S.C., Davis, K.L., Amir, E.D., Tadmor, M.D., Litvin, O., Fienberg, H.G., Jager, A., Zunder, E.R., Finck, R., Gedman, A.L., Radtke, I., Downing, J.R., Pe’er, D., Nolan, G.P., 2015. Data-Driven Phenotypic Dissection of AML Reveals Progenitor-like Cells that Correlate with Prognosis. Cell 162, 184–197. https://doi.org/10.1016/j.cell.2015.05.047

Levitin, H.M., Yuan, J., Cheng, Y.L., Ruiz, F.J., Bush, E.C., Bruce, J.N., Canoll, P., Iavarone, A., Lasorella, A., Blei, D.M., Sims, P.A., 2019. De novo gene signature identification from single-cell RNA-seq with hierarchical Poisson factorization. Mol Syst Biol 15, e8557. https://doi.org/10.15252/msb.20188557

Lindestam Arlehamn, C.S., Dhanwani, R., Pham, J., Kuan, R., Frazier, A., Rezende Dutra, J., Phillips, E., Mallal, S., Roederer, M., Marder, K.S., Amara, A.W., Standaert, D.G., Goldman, J.G., Litvan, I., Peters, B., Sulzer, D., Sette, A., 2020. α-Synuclein-specific T cell reactivity is associated with preclinical and early Parkinson’s disease. Nat Commun 11, 1875. https://doi.org/10.1038/s41467-020-15626-w

Lodygin, D., Hermann, M., Schweingruber, N., Flügel-Koch, C., Watanabe, T., Schlosser, C., Merlini, A., Körner, H., Chang, H.-F., Fischer, H.J., Reichardt, H.M., Zagrebelsky, M., Mollenhauer, B., Kügler, S., Fitzner, D., Frahm, J., Stadelmann, C., Haberl, M., Odoardi, F., Flügel, A., 2019. β-Synuclein-reactive T cells induce autoimmune CNS grey matter degeneration. Nature 566, 503–508. https://doi.org/10.1038/s41586-019-0964-2

Love, M.I., Huber, W., Anders, S., 2014. Moderated estimation of fold change and dispersion for RNA-seq data with DESeq2. Genome Biology 15, 550. https://doi.org/10.1186/s13059-014-0550-8

Lun, A.T.L., Bach, K., Marioni, J.C., 2016. Pooling across cells to normalize single-cell RNA sequencing data with many zero counts. Genome Biol 17, 75. https://doi.org/10.1186/s13059-016-0947-7

Matsuda, W., Furuta, T., Nakamura, K.C., Hioki, H., Fujiyama, F., Arai, R., Kaneko, T., 2009. Single nigrostriatal dopaminergic neurons form widely spread and highly dense axonal arborizations in the neostriatum. J. Neurosci. 29, 444–453. https://doi.org/10.1523/JNEUROSCI.4029-08.2009

McDole, J.R., Danzer, S.C., Pun, R.Y.K., Chen, Y., Johnson, H.L., Pirko, I., Johnson, A.J., 2010. Rapid Formation of Extended Processes and Engagement of Theiler’s Virus-Infected Neurons by CNS-Infiltrating CD8 T Cells. The American Journal of Pathology 177, 1823–1833. https://doi.org/10.2353/ajpath.2010.100231

McGeer, P.L., Itagaki, S., Boyes, B.E., McGeer, E.G., 1988. Reactive microglia are positive for HLA-DR in the substantia nigra of Parkinson’s and Alzheimer’s disease brains. Neurology 38, 1285–1291. https://doi.org/10.1212/wnl.38.8.1285

Min, W., Pober, J.S., Johnson, D.R., 1996. Kinetically coordinated induction of TAP1 and HLA class I by IFN-gamma: the rapid induction of TAP1 by IFN-gamma is mediated by Stat1 alpha. J Immunol 156, 3174–3183.

Mo, M.-S., Li, G.-H., Sun, C.-C., Huang, S.-X., Wei, L., Zhang, L.-M., Zhou, M.-M., Wu, Z.-H., Guo, W.-Y., Yang, X.-L., Chen, C.-J., Qu, S.-G., He, J.-X., Xu, P.-Y., 2018. Dopaminergic neurons show increased low-molecular-mass protein 7 activity induced by 6-hydroxydopamine in vitro and in vivo. Translational Neurodegeneration 7, 19. https://doi.org/10.1186/s40035-018-0125-9

Mogi, M., Kondo, T., Mizuno, Y., Nagatsu, T., 2007. p53 protein, interferon-gamma, and NF-kappaB levels are elevated in the parkinsonian brain. Neurosci Lett 414, 94–97. https://doi.org/10.1016/j.neulet.2006.12.003

Morales, J., Homey, B., Vicari, A.P., Hudak, S., Oldham, E., Hedrick, J., Orozco, R., Copeland, N.G., Jenkins, N.A., McEvoy, L.M., Zlotnik, A., 1999. CTACK, a skin-associated chemokine that preferentially attracts skin-homing memory T cells. Proc Natl Acad Sci U S A 96, 14470–14475. https://doi.org/10.1073/pnas.96.25.14470

Mount, M.P., Lira, A., Grimes, D., Smith, P.D., Faucher, S., Slack, R., Anisman, H., Hayley, S., Park, D.S., 2007. Involvement of interferon-gamma in microglial-mediated loss of dopaminergic neurons. J. Neurosci. 27, 3328–3337. https://doi.org/10.1523/JNEUROSCI.5321-06.2007

Müllner, D., 2013. fastcluster: Fast Hierarchical, Agglomerative Clustering Routines for R and Python. Journal of Statistical Software 53, 1–18. https://doi.org/10.18637/jss.v053.i09

Naik, J., Hau, C.M., Bloemendaal, L. ten, Mok, K.S., Hajji, N., Wehman, A.M., Meisner, S., Muncan, V., Paauw, N.J., Vries, H.E. de, Nieuwland, R., Paulusma, C.C., Bosma, P.J., 2019. The P4-ATPase ATP9A is a novel determinant of exosome release. PLOS ONE 14, e0213069. https://doi.org/10.1371/journal.pone.0213069

Namiki, S., Nakamura, T., Oshima, S., Yamazaki, M., Sekine, Y., Tsuchiya, K., Okamoto, R., Kanai, T., Watanabe, M., 2005. IRF-1 mediates upregulation of LMP7 by IFN-gamma and concerted expression of immunosubunits of the proteasome. FEBS Lett 579, 2781–2787. https://doi.org/10.1016/j.febslet.2005.04.012

Neumann, H., Schmidt, H., Cavalié, A., Jenne, D., Wekerle, H., 1997. Major histocompatibility complex (MHC) class I gene expression in single neurons of the central nervous system: differential regulation by interferon (IFN)-gamma and tumor necrosis factor (TNF)-alpha. J Exp Med 185, 305–316. https://doi.org/10.1084/jem.185.2.305

Popko, B., Corbin, J.G., Baerwald, K.D., Dupree, J., Garcia, A.M., 1997. The effects of interferon-gamma on the central nervous system. Mol. Neurobiol. 14, 19–35. https://doi.org/10.1007/BF02740619

Poulin, J.-F., Zou, J., Drouin-Ouellet, J., Kim, K.-Y.A., Cicchetti, F., Awatramani, R.B., 2014. Defining midbrain dopaminergic neuron diversity by single-cell gene expression profiling. Cell Rep 9, 930–943. https://doi.org/10.1016/j.celrep.2014.10.008

Rall, G.F., 1998. CNS Neurons: The Basis and Benefits of Low Class I Major Histocompatibility Complex Expression, in: Whitton, J.L. (Ed.), Antigen Presentation, Current Topics in Microbiology and Immunology. Springer, Berlin, Heidelberg, pp. 115–134. https://doi.org/10.1007/978-3-642-72045-1_6

Rico-Bautista, E., Flores-Morales, A., Fernández-Pérez, L., 2006. Suppressor of cytokine signaling (SOCS) 2, a protein with multiple functions. Cytokine & Growth Factor Reviews 17, 431–439. https://doi.org/10.1016/j.cytogfr.2006.09.008

Robertson, I.B., Horiguchi, M., Zilberberg, L., Dabovic, B., Hadjiolova, K., Rifkin, D.B., 2015. Latent TGF-β-binding proteins. Matrix Biol 47, 44–53. https://doi.org/10.1016/j.matbio.2015.05.005

Rose, R.W., Vorobyeva, A.G., Skipworth, J.D., Nicolas, E., Rall, G.F., 2007. Altered Levels of STAT1 and STAT3 Influence the Neuronal Response to Interferon Gamma. J Neuroimmunol 192, 145–156. https://doi.org/10.1016/j.jneuroim.2007.10.007

Rostami, J., Fotaki, G., Sirois, J., Mzezewa, R., Bergström, J., Essand, M., Healy, L., Erlandsson, A., 2020. Astrocytes have the capacity to act as antigen-presenting cells in the Parkinson’s disease brain. Journal of Neuroinflammation 17, 119. https://doi.org/10.1186/s12974-020-01776-7

Sadakata, T., Mizoguchi, A., Sato, Y., Katoh-Semba, R., Fukuda, M., Mikoshiba, K., Furuichi, T., 2004. The Secretory Granule-Associated Protein CAPS2 Regulates Neurotrophin Release and Cell Survival. J Neurosci 24, 43–52. https://doi.org/10.1523/JNEUROSCI.2528-03.2004

Samarajiwa, S.A., Forster, S., Auchettl, K., Hertzog, P.J., 2009. INTERFEROME: the database of interferon regulated genes. Nucleic Acids Res 37, D852–857. https://doi.org/10.1093/nar/gkn732

Santos, S.E.D., Medeiros, M., Porfirio, J., Tavares, W., Pessôa, L., Grinberg, L., Leite, R.E.P., Ferretti-Rebustini, R.E.L., Suemoto, C.K., Filho, W.J., Noctor, S.C., Sherwood, C.C., Kaas, J.H., Manger, P.R., Herculano-Houzel, S., 2020. Similar Microglial Cell Densities across Brain Structures and Mammalian Species: Implications for Brain Tissue Function. J. Neurosci. 40, 4622–4643. https://doi.org/10.1523/JNEUROSCI.2339-19.2020

Saunders, A., Macosko, E., Wysoker, A., Goldman, M., Krienen, F., de Rivera, H., Bien, E., Baum, M., Wang, S., Goeva, A., Nemesh, J., Kamitaki, N., Brumbaugh, S., Kulp, D., McCarroll, S.A., 2018. Molecular Diversity and Specializations among the Cells of the Adult Mouse Brain. Cell 174, 1015-1030.e16. https://doi.org/10.1016/j.cell.2018.07.028

Schroder, K., Hertzog, P.J., Ravasi, T., Hume, D.A., 2004. Interferon-γ: an overview of signals, mechanisms and functions. Journal of Leukocyte Biology 75, 163–189. https://doi.org/10.1189/jlb.0603252

Shekhar, K., Lapan, S.W., Whitney, I.E., Tran, N.M., Macosko, E.Z., Kowalczyk, M., Adiconis, X., Levin, J.Z., Nemesh, J., Goldman, M., McCarroll, S.A., Cepko, C.L., Regev, A., Sanes, J.R., 2016. Comprehensive Classification of Retinal Bipolar Neurons by Single-Cell Transcriptomics. Cell 166, 1308-1323.e30. https://doi.org/10.1016/j.cell.2016.07.054

Shuai, K., Stark, G.R., Kerr, I.M., Darnell, J.E., 1993. A single phosphotyrosine residue of Stat91 required for gene activation by interferon-gamma. Science 261, 1744–1746. https://doi.org/10.1126/science.7690989

Snyder, M.E., Finlayson, M.O., Connors, T.J., Dogra, P., Senda, T., Bush, E., Carpenter, D., Marboe, C., Benvenuto, L., Shah, L., Robbins, H., Hook, J.L., Sykes, M., D’Ovidio, F., Bacchetta, M., Sonett, J.R., Lederer, D.J., Arcasoy, S., Sims, P.A., Farber, D.L., 2019. Generation and persistence of human tissue-resident memory T cells in lung transplantation. Sci Immunol 4. https://doi.org/10.1126/sciimmunol.aav5581

Sommer, A., Marxreiter, F., Krach, F., Fadler, T., Grosch, J., Maroni, M., Graef, D., Eberhardt, E., Riemenschneider, M.J., Yeo, G.W., Kohl, Z., Xiang, W., Gage, F.H., Winkler, J., Prots, I., Winner, B., 2018. Th17 Lymphocytes Induce Neuronal Cell Death in a Human iPSC-Based Model of Parkinson’s Disease. Cell Stem Cell 23, 123-131.e6. https://doi.org/10.1016/j.stem.2018.06.015

Strickland, M.R., Koller, E.J., Deng, D.Z., Ceballos-Diaz, C., Golde, T.E., Chakrabarty, P., 2018. Ifngr1 and Stat1 mediated canonical Ifn-γ signaling drives nigrostriatal degeneration. Neurobiol. Dis. 110, 133–141. https://doi.org/10.1016/j.nbd.2017.11.007

Stüve, O., Youssef, S., Slavin, A.J., King, C.L., Patarroyo, J.C., Hirschberg, D.L., Brickey, W.J., Soos, J.M., Piskurich, J.F., Chapman, H.A., Zamvil, S.S., 2002. The Role of the MHC Class II Transactivator in Class II Expression and Antigen Presentation by Astrocytes and in Susceptibility to Central Nervous System Autoimmune Disease. The Journal of Immunology 169, 6720–6732. https://doi.org/10.4049/jimmunol.169.12.6720

Sulzer, D., Alcalay, R.N., Garretti, F., Cote, L., Kanter, E., Agin-Liebes, J., Liong, C., McMurtrey, C., Hildebrand, W.H., Mao, X., Dawson, V.L., Dawson, T.M., Oseroff, C., Pham, J., Sidney, J., Dillon, M.B., Carpenter, C., Weiskopf, D., Phillips, E., Mallal, S., Peters, B., Frazier, A., Lindestam Arlehamn, C.S., Sette, A., 2017. T cells from patients with Parkinson’s disease recognize α-synuclein peptides. Nature 546, 656–661. https://doi.org/10.1038/nature22815

Svensson, M., Marsal, J., Ericsson, A., Carramolino, L., Brodén, T., Márquez, G., Agace, W.W., 2002. CCL25 mediates the localization of recently activated CD8alphabeta(+) lymphocytes to the small-intestinal mucosa. J Clin Invest 110, 1113–1121. https://doi.org/10.1172/JCI15988

Tanaka, Y., Ono, N., Shima, T., Tanaka, G., Katoh, Y., Nakayama, K., Takatsu, H., Shin, H.-W., 2016. The phospholipid flippase ATP9A is required for the recycling pathway from the endosomes to the plasma membrane. Mol Biol Cell 27, 3883–3893. https://doi.org/10.1091/mbc.E16-08-0586

Taylor, G.A., 2007. IRG proteins: key mediators of interferon-regulated host resistance to intracellular pathogens. Cell Microbiol 9, 1099–1107. https://doi.org/10.1111/j.1462-5822.2007.00916.x

Ugras, S., Daniels, M.J., Fazelinia, H., Gould, N.S., Yocum, A.K., Luk, K.C., Luna, E., Ding, H., McKennan, C., Seeholzer, S., Martinez, D., Evans, P., Brown, D., Duda, J.E., Ischiropoulos, H., 2018. Induction of the Immunoproteasome Subunit Lmp7 Links Proteostasis and Immunity in α-Synuclein Aggregation Disorders. EBioMedicine 31, 307–319. https://doi.org/10.1016/j.ebiom.2018.05.007

van den Elsen, P.J., Holling, T.M., Kuipers, H.F., van der Stoep, N., 2004. Transcriptional regulation of antigen presentation. Current Opinion in Immunology 16, 67–75. https://doi.org/10.1016/j.coi.2003.11.015

Vass, K., Lassmann, H., 1990. Intrathecal application of interferon gamma. Progressive appearance of MHC antigens within the rat nervous system. Am J Pathol 137, 789–800.

Wang, F., Flanagan, J., Su, N., Wang, L.-C., Bui, S., Nielson, A., Wu, X., Vo, H.-T., Ma, X.-J., Luo, Y., 2012. RNAscope: A Novel in Situ RNA Analysis Platform for Formalin-Fixed, Paraffin-Embedded Tissues. The Journal of Molecular Diagnostics 14, 22–29. https://doi.org/10.1016/j.jmoldx.2011.08.002

Xie, Z., Bailey, A., Kuleshov, M.V., Clarke, D.J.B., Evangelista, J.E., Jenkins, S.L., Lachmann, A., Wojciechowicz, M.L., Kropiwnicki, E., Jagodnik, K.M., Jeon, M., Ma’ayan, A., 2021. Gene Set Knowledge Discovery with Enrichr. Current Protocols 1, e90. https://doi.org/10.1002/cpz1.90

Yshii, L.M., Gebauer, C.M., Pignolet, B., Mauré, E., Quériault, C., Pierau, M., Saito, H., Suzuki, N., Brunner-Weinzierl, M., Bauer, J., Liblau, R., 2016. CTLA4 blockade elicits paraneoplastic neurological disease in a mouse model. Brain 139, 2923–2934. https://doi.org/10.1093/brain/aww225

Yuan, J., Levitin, H.M., Frattini, V., Bush, E.C., Boyett, D.M., Samanamud, J., Ceccarelli, M., Dovas, A., Zanazzi, G., Canoll, P., Bruce, J.N., Lasorella, A., Iavarone, A., Sims, P.A., 2018. Single-cell transcriptome analysis of lineage diversity in high-grade glioma. Genome Medicine 10, 57. https://doi.org/10.1186/s13073-018-0567-9

Zhao, W., Dovas, A., Spinazzi, E.F., Levitin, H.M., Banu, M.A., Upadhyayula, P., Sudhakar, T., Marie, T., Otten, M.L., Sisti, M.B., Bruce, J.N., Canoll, P., Sims, P.A., 2021. Deconvolution of cell type-specific drug responses in human tumor tissue with single-cell RNA-seq. Genome Medicine 13, 82. https://doi.org/10.1186/s13073-021-00894-y

Zhou, F., 2009. Molecular mechanisms of IFN-gamma to up-regulate MHC class I antigen processing and presentation. Int Rev Immunol 28, 239–260. https://doi.org/10.1080/08830180902978120

