## Supplementary Figures for "Conserved and cell type-specific transcriptional responses to IFN-γ in the ventral midbrain"

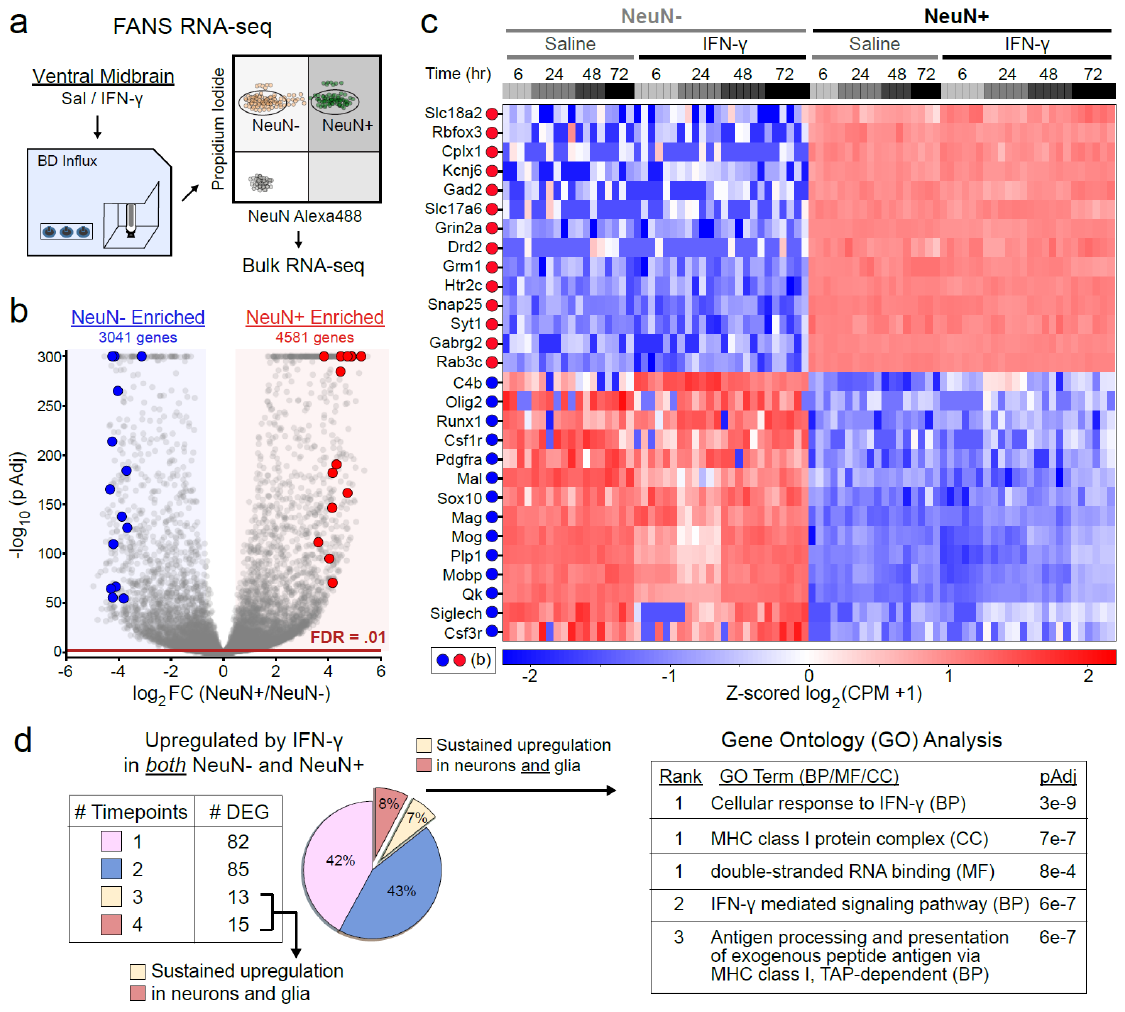

**Supplementary Figure 1: Purity of nuclear sorting and sustained upregulation of canonical IFN-γ genes in both neurons and glia**

(**a**) Fluorescence-activated nuclear sorting (FANS) RNA-seq schematic. Nuclei are gated on propidium iodide and anti-NeuN fluorescence to sort NeuN- and NeuN+ populations. Bulk RNA-seq is conducted in 96 well-plates on ~1000 sorted nuclei per well.

(**b**) Volcano plot for NeuN+ vs. NeuN- sorted nuclei (n = 36 NeuN-, n = 48 NeuN+). Enriched gene numbers are based on |Log2FC| > 0.5 and pAdj < 0.01 (*DESeq2* Wald test). Red and blue points are shown in the heatmap in (c).

(**c**) Heatmap of z-scored mRNA abundances, normalized as log2(CPM + 1). Each column represents a bulk RNA-seq replicate of ~1000 sorted nuclei (n = 4-6 replicates from 2-3 mice for each treatment/timepoint). Neuronal and glial genes are red/blue in (b).

(**d**) *Left*: For all genes upregulated in *both* NeuN- and NeuN+ samples at any timepoint, the number (*left*) and fraction (*right*) upregulated at 1-4 timepoints is shown (e.g., 82 mRNAs are upregulated only at one timepoint, while 15 mRNAs are upregulated at all four timepoints). *Right*: GO analysis for the 28 mRNAs upregulated at ≥ 3 timepoints in both NeuN- and NeuN+ samples from left.

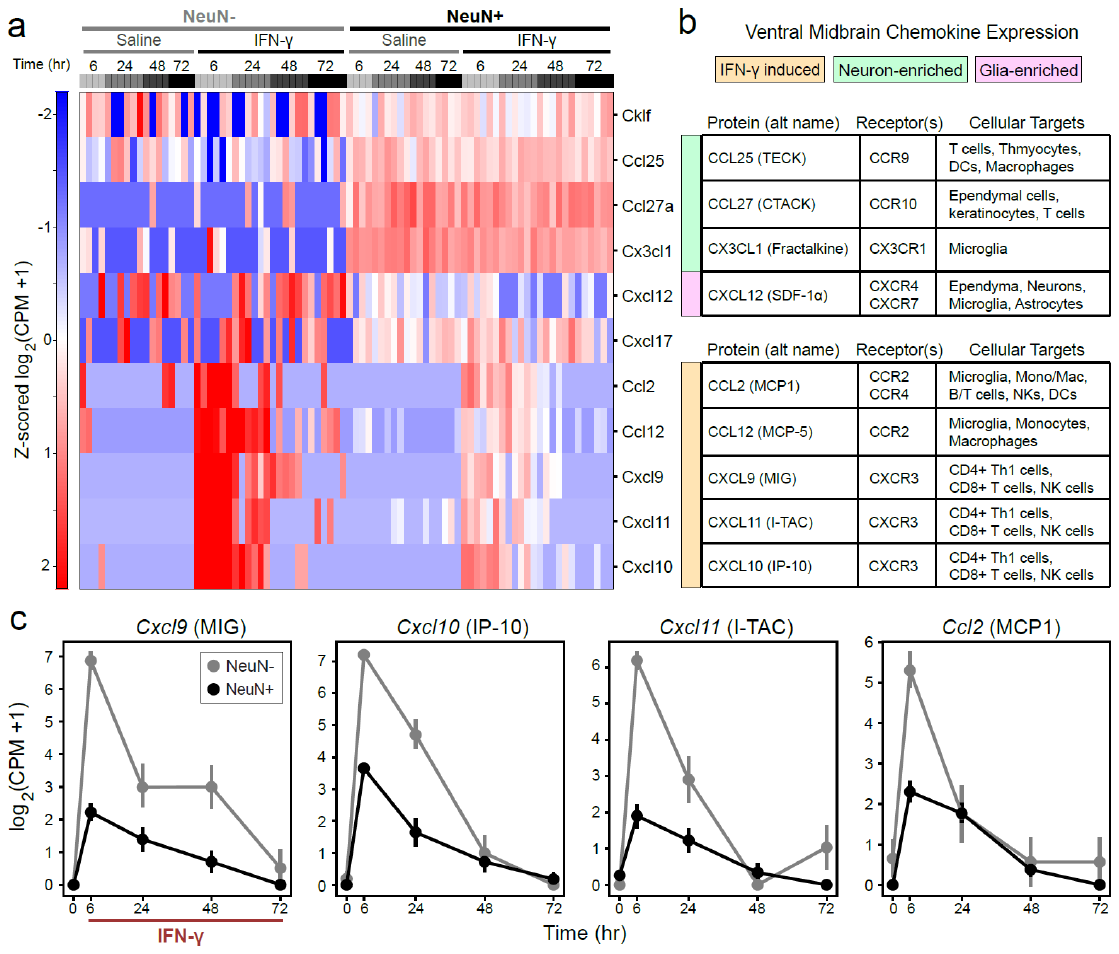

**Supplementary Figure 2: Chemokine expression induced by IFN-γ**

(**a**) Heatmap of z-scored mRNA abundances, normalized as log2(CPM + 1). Each column represents a bulk RNA-seq replicate of ~1000 sorted nuclei (n = 4-6 replicates from 2-3 mice for each treatment/timepoint).

(**b**) Table summary of encoded chemokines from (a), their receptors, and cellular targets.

(**c**) Mean ± SEM for mRNA abundance in NeuN-/NeuN+ samples, normalized as log2(CPM + 1). Samples as in (a), (n = 4-6 replicates each of NeuN-/+ nuclei from 2-3 mice for each treatment/timepoint). Saline samples from all timepoints are together at t=0.

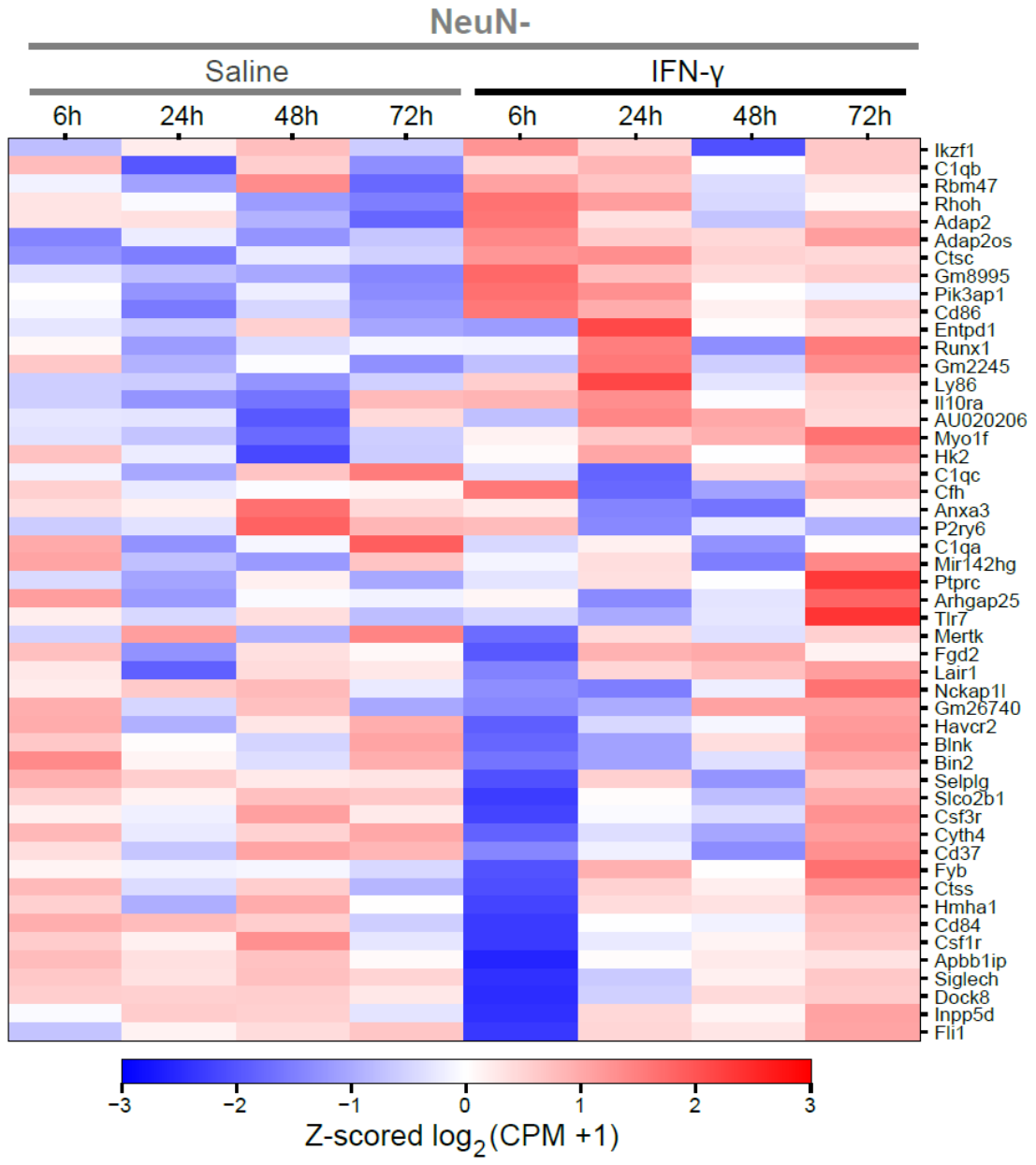

**Supplementary Figure 3: Acute differential regulation of microglial lineage markers by IFN-γ**

Heatmap of mRNA abundances for the top 50 microglial marker genes (genes identified via snRNA-seq data, see **Figure 3**). Each column represents the z-score of the average log2(CPM + 1) for the indicated treatment group (n = 4-6 replicates of ~1000 sorted nuclei from 2-3 mice for each treatment/timepoint).

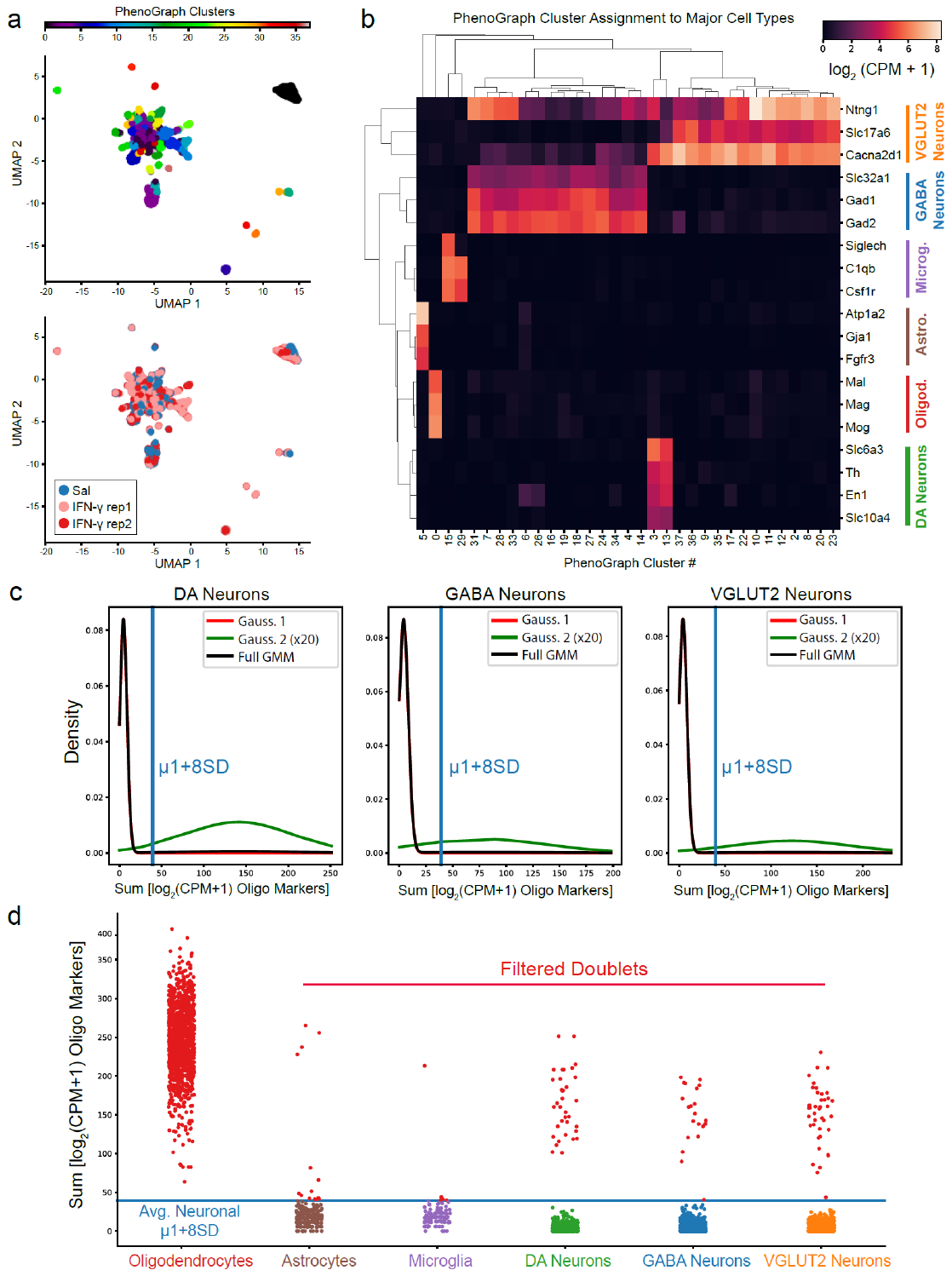
 **Supplementary Figure 4: Assignment of PhenoGraph clusters to six major cell types and filtering of oligodendrocyte doublets**

(**a**) UMAP embedding of single nucleus RNA-seq profiles (5,464) from the ventral midbrain. *Upper*: Complete set of PhenoGraph clusters before collapse to six major cell types. *Lower*: Mouse sample origin (n = 1 for saline, n = 2 for IFN-γ).

(**b**) Clustered heatmap of the average expression (log2[CPM + 1]) of the indicated genes within each PhenoGraph cluster. PhenoGraph clusters were assigned to the six major cell types indicated on the right based on hierarchical clustering (top).

(**c**) Gaussian mixture models for unfiltered DA, GABA, or VGLUT2 neurons. The black trace indicates the fitted GMM; the red trace indicates component 1 (single neurons); the green trace indicates component 2 (multiplets), scaled up by a factor of 20 for visualization. The mean of the first component plus eight standard deviations is shown in blue.

(**d**) Filtering of oligodendrocyte doublets from other major cell types. The blue trace indicates the average neuronal component 1 mean plus eight standard deviations, indicated in the GMM’s above. Filtered doublets are indicated in red. The same procedure was repeated to remove astrocyte- and microglia-containing doublets (not shown).

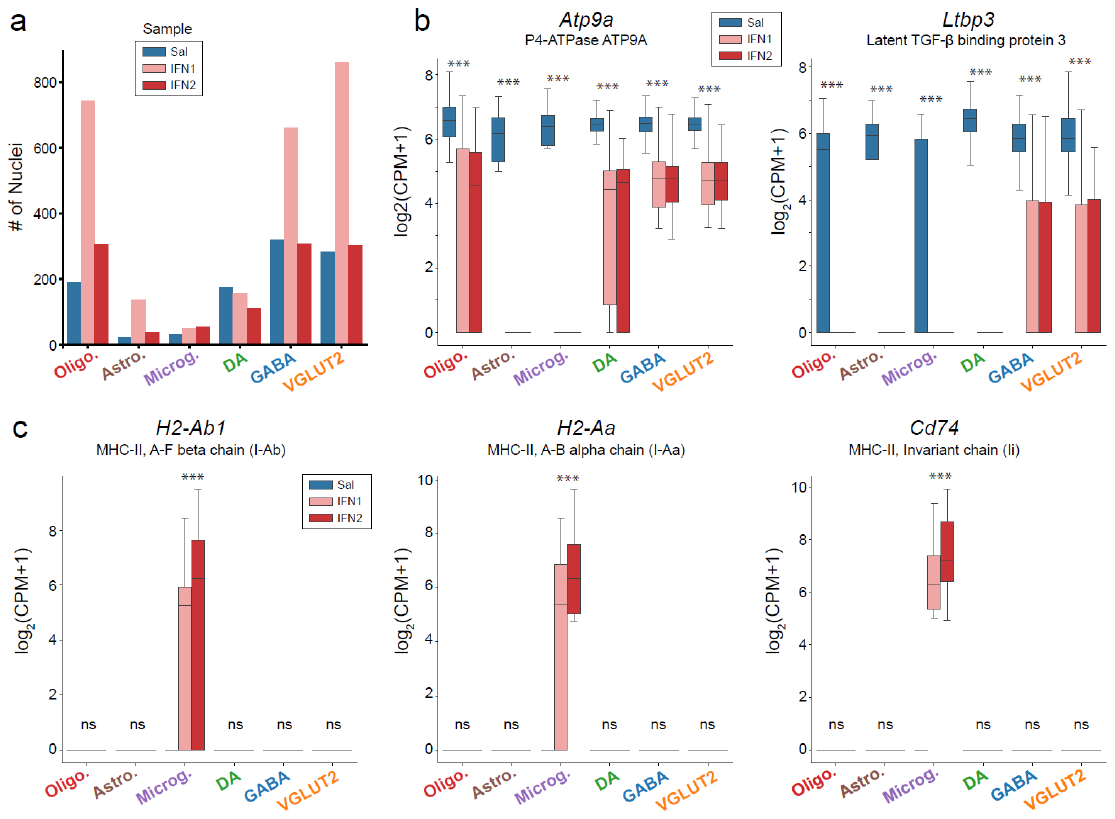

**Supplementary Figure 5: Nuclei distribution per mouse, cell type-independent downregulation of Atp9a and Ltbp3 by IFN-γ, and microglia-specific MHC-II expression**

(**a**) Number of nuclei from each mouse sample within the major cell type groups.

(**b**) RNA expression (log2 [CPM + 1]) of *Atp9a* and *Ltbp3* in each major cell type. Both are significantly downregulated by IFN-γ in all cell types; *** q < 0.05, modified Mann-Whitney (see **Methods**).

(**c**) RNA expression (log2 [CPM + 1]) of MHC-II genes *H2-Ab1*, *H2-Aa*, and *Cd74* in each major cell type; upregulation by IFN-γ occurs only in microglia; *** q < 0.05, Mann-Whitney (see **Methods**).

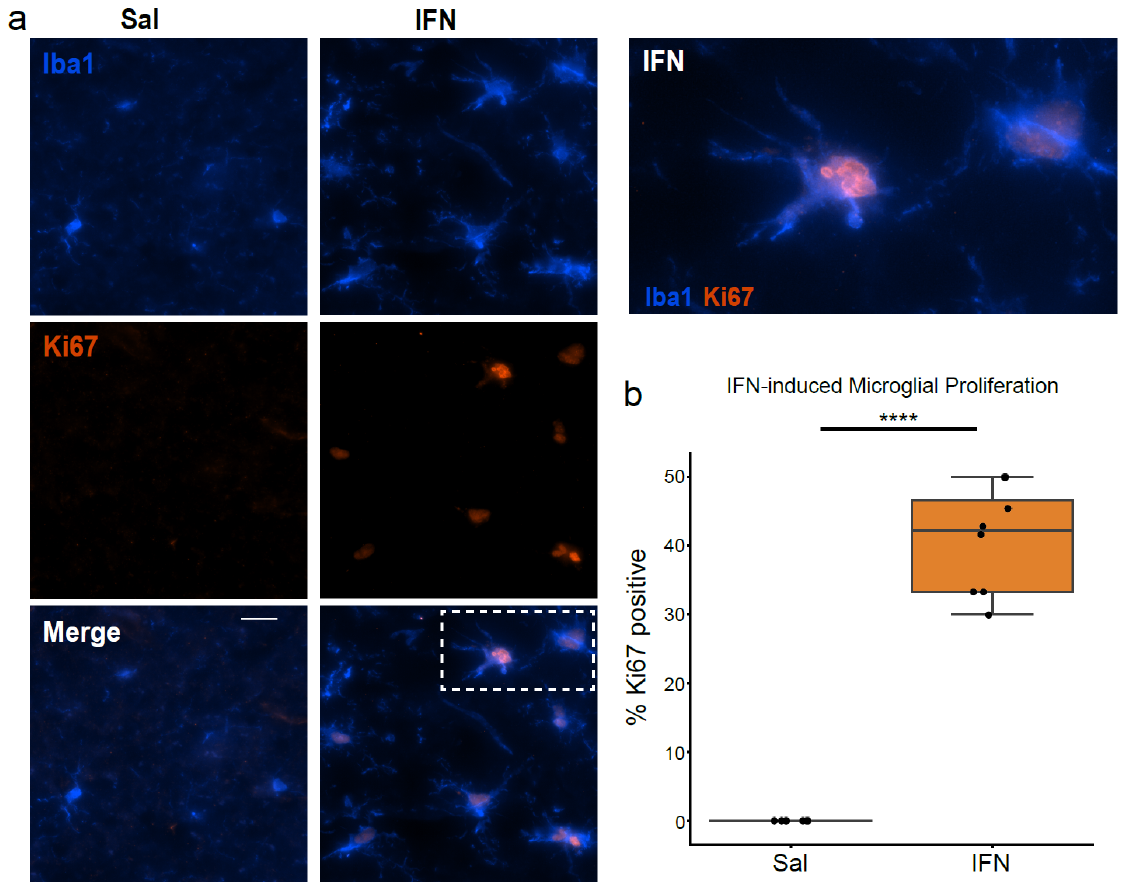

**Supplementary Figure 6: IFN-γ induces microglial proliferation**

(**a**) Representative images of Iba1^+^ microglia in the substantia nigra, which express the proliferation marker Ki67 following exposure to IFN-γ. Scale bar: 20 µm.

(**b**) Quantification of Ki67^+^/Iba1^+^ cells as a percentage of all Iba1+ cells in the substantia nigra, related to (a). **** p < .001, Mann-Whitney U test, 2-3 fields from n = 3 mice for each treatment.

**Supplementary Figure 7: Sub-clustering of dopamine neurons**

(**a**) Heatmap of normalized expression, log_2_(counts per 10,000 + 1), for the top markers of each subcluster.

(**b**) UMAP embedding of dopamine neuronal subclusters with overlay of cluster ID, *Slc6a3*/DAT expression, or normalized expression, log_2_(CPM+1), for selected marker genes.

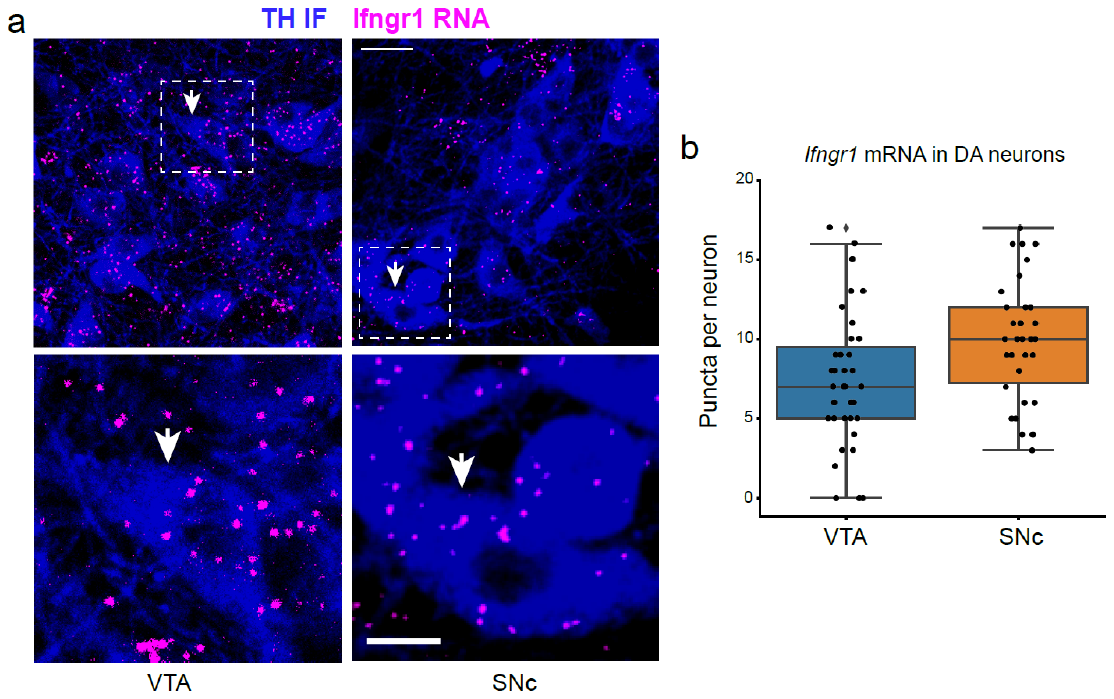

**Supplementary Figure 8: *Ifngr1* mRNA expression in mDA neurons**

(**a**) Ifngr1 RNA FISH in the substantia nigra pars compacta (SNc) or ventral tegmental area (VTA). White arrowheads indicate TH^+^ mDA neurons in the inset (*lower*). Scale bars, upper: 25 µm, lower: 10 µm.

(**b**) Quantification of *Ifngr1* mRNA puncta in mDA neurons, related to (a). 10-15 neurons were quantified per region per mouse, from n=3 mice.

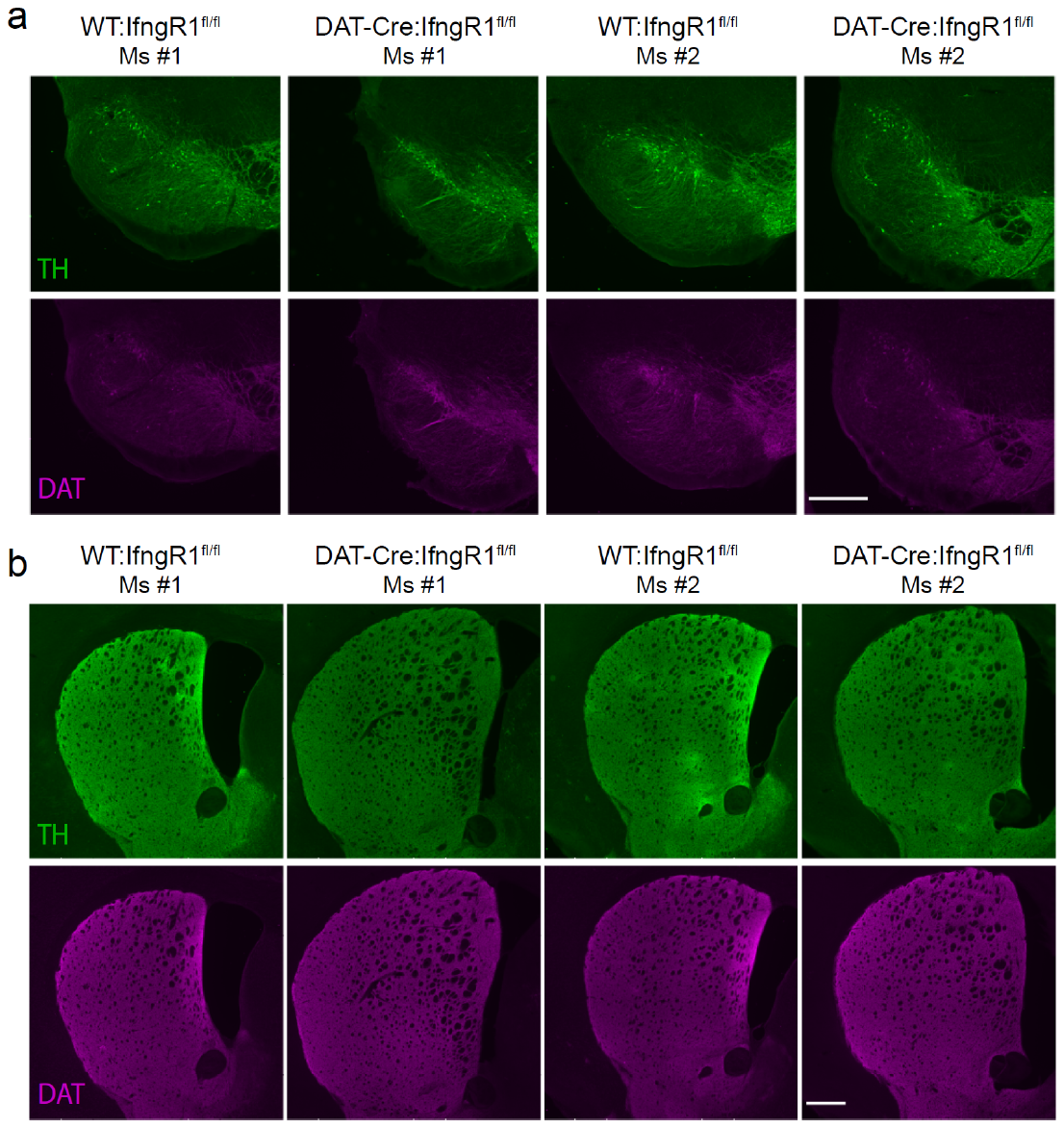

**Supplementary Figure 9: No gross abnormalities in DA neuronal development and striatal innervation in DAT^IRES-^Cre:Ifngr1^fl/fl^ mice**

(**a-b**) TH and DAT immunofluorescence in the substantia nigra (a) or striatum (b) of WT:Ifngr1^fl/fl^ or DAT-Cre:Ifngr1^fl/fl^ mice. Scale bars: 500 µm.
